# Branched DNA Processing by a Thermostable CAS-Cas4 from *Thermococcus onnurineus*: Expanding Biochemical Landscape of Nuclease Activity

**DOI:** 10.1101/2024.11.29.625776

**Authors:** Muskan Jain, Asish Kumar Pattnayak, Sakshi Aggarwal, Praveen Rai, Kavya J, Sanjeev Chandrayan, Manisha Goel, Vineet Gaur

**Author notes:** Corresponding authors: **Dr. Vineet Gaur:** National Institute of Plant Genome Research, Aruna Asaf Ali Marg, New Delhi 110067, India., **Dr. Manisha Goel:** Department of Biophysics, University of Delhi South Campus, Benito Juarez Road, New Delhi, Delhi 110021, India.

## Abstract

The adaptive immune function of CRISPR-Cas systems in bacteria and archaea is mediated through **<u>C</u>**RISPR-**<u>A</u>**ssociated Proteins (Cas). The adaptation module, typically involving Cas1, Cas2, and Cas4, helps integrate viral “spacer” sequences into the host genome. Cas4 proteins are classified into two types based on neighboring genes: CAS-Cas4, flanked by other *cas* genes, and Solo-Cas4, which exist independently. While CAS-Cas4 proteins are implicated in adaptation, they remain biochemically uncharacterized in archaea, unlike archaeal Solo-Cas4 proteins. This study biochemically characterizes TON_0321, a CAS-Cas4 protein from the Type IV-C CRISPR cassette of *Thermococcus onnurineus*. TON_0321 exhibits 5′ to 3′ exonuclease activity and unique structure-dependent endonuclease activity, shedding light on CAS-Cas4 functional diversity. A distinct spatial organization of the catalytic site, angled with the positively charged patch on the protein surface, enables TON_0321 to recognize branching points in DNA substrates. Furthermore, this spatial arrangement facilitates cleavage 2 to 3 nucleotides away from the branch point in the 5′ direction, demonstrating structure-specific endonuclease activity.

## Introduction

**<u>C</u>**lustered **<u>R</u>**egularly **<u>I</u>**nterspaced **<u>S</u>**hort **<u>P</u>**alindromic **R**epeats (CRISPR) are a type of acquired immune system in bacteria and archaea (1–3), which are already proving to be powerful tools in targeted genome editing (1–9). Their function is mediated by a variety of proteins, collectively termed **<u>C</u>**RISPR-**<u>as</u>**sociated proteins (Cas proteins). Currently, CRISPR function is understood to be mediated by either a group of proteins (class 1) or via a single protein with multiple functional domains working as an effector protein (class 2). These two classes are further subdivided into several types and subtypes. The classification is based on the presence of “unique signature” proteins to each CRISPR-Cas type. The class 1 systems encompass type I, III, and IV systems, whereas the class 2 CRISPR systems are subdivided into three types: II, V, and VI involving Cas9, Cas12, and Cas13 proteins, respectively. Despite the diverse set of proteins serving as effectors among different CRISPR types, all CRISPR-Cas systems appear to operate in three stages: adaptation, where foreign DNA is incorporated into CRISPR arrays; expression, involving transcription and processing of CRISPR RNA (crRNA) from integrated spacers; and interference, where crRNA is used to guide the degradation of the invading genomic material, conferring immunity against re-invading pathogen (10–14). The adaptation modules of CRISPR systems, typically contain Cas1 and Cas2 which excise, process, and integrate foreign DNA (pre-spacers) into the CRISPR-Cas loci as a new spacer (15, 16). However, several CRIPSR types (I, II, and V) have an additional protein, termed Cas4, involved in the adaptation step (15) (Fig. 1A). Processing pre-spacers into spacer fragments of the correct size with appropriate ends is quintessential for integration into a specific orientation (17–19). Incorrect spacer processing compromises the defense response (20, 21). Cas4 proteins have been shown to determine the length and orientation of the new spacers by recognizing specific short sequences called **<u>p</u>**rotospacer **<u>a</u>**djacent **<u>m</u>**otifs (PAMs) (20, 21).

**Figure 1.**
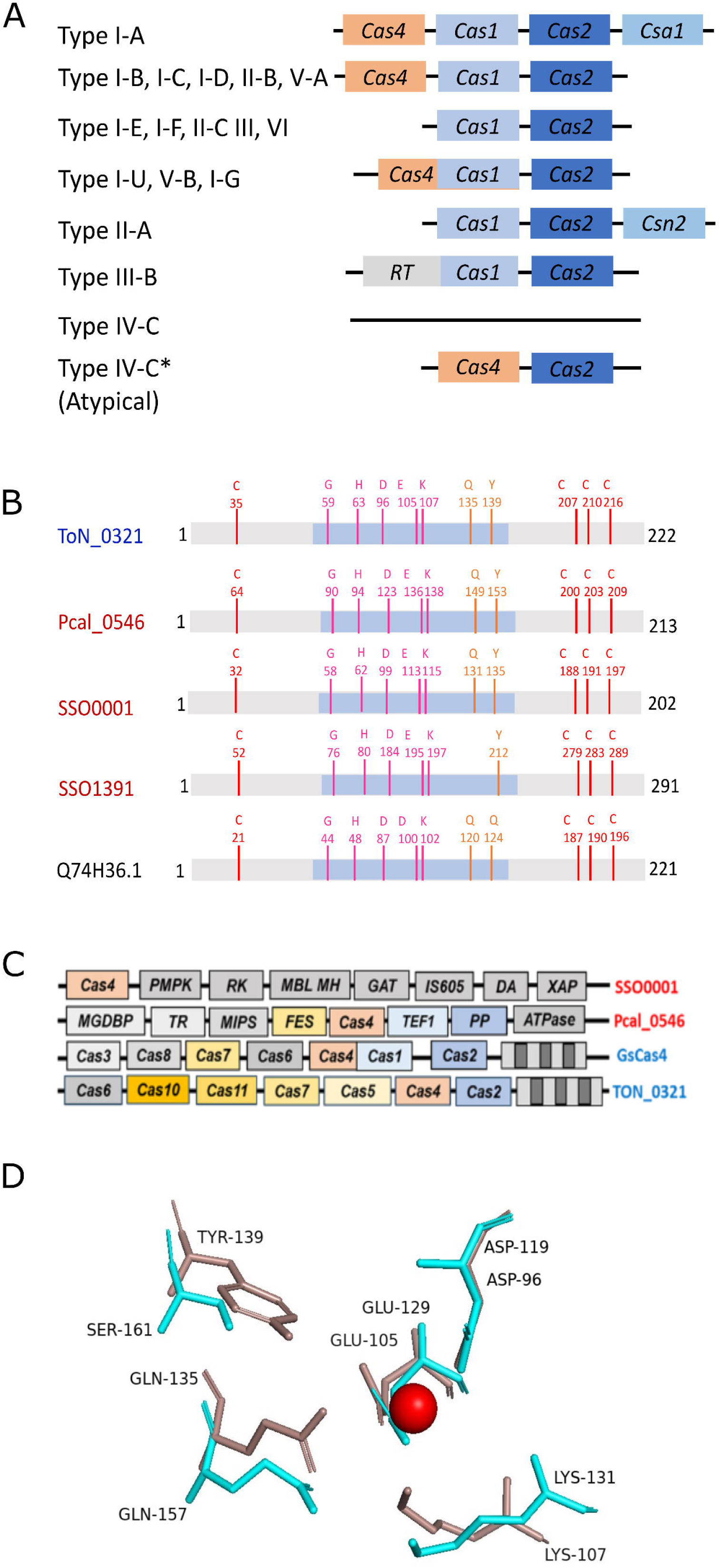
Diversity among Cas4 proteins. **(A)** Block diagram showing the typical organization of adaptation module in different CRISPR subtypes. Adaptation module usually consists of Cas1, Cas2 and Cas4 proteins. It may contain other proteins in specific subtypes. A few of the CRISPR subtypes does not include Cas4. The typical type IV-C CRISPR subtype does not contain adaptation module. The atypical type IV-C CRISPR cassette from *Thermococcus onnurineus NA1* shows the presence of Cas2 and Cas4 adaptation proteins. Type I-A, I-B…III-B: different CRISPR subtypes; Type IV-C: typical type IV-C cassette; Type IV-C*: atypical case of *Thermococcus onnurineus NA1.* **(B)** A cartoon depicting conserved amino acid residues among various characterized Cas4 proteins: TON_0321 (*Thermococcus onnurineus NA1),* Pcal_0546 (*Pyrobaculum calidifontis*), SSO0001 (*Sulfolobus solfataricus*), SSO1391 (*Sulfolobus solfataricus*), and GsCas4 (*Geobacter sulfurreducens*). Four cysteines involved in chelating Fe-S cluster are shown in red. RecB active site residues are shown in pink and orange; the blue-shaded region depicts the RecB-like domain of Cas4 proteins. **(C)** A schematic showing the difference in gene cassettes of SSO0001 from *Sulfolobus solfataricus*, Pcal_0546 from *Pyrobaculum calidifontis*, GsCas4 from *Geobacter sulfurreducens* and TON_0321 from *Thermococcus onnurineus* NA1). **(D)** Comparison of the catalytic sites of λ exonuclease (cyan; PDB: 1AVQ) and TON_0321 (red, model) depicting the conservation of active site residues. The metal ion at the active site is shown as a sphere.

Cas4 proteins belong to the PD-(D/E)XK endonuclease-like domain superfamily (22). PD-(D/E)XK nucleases form a highly diverse superfamily of enzymes comprising of restriction endonucleases (*e.g.,* EcoRI, XhoI) (23), resolvases (*e.g.,* phage lambda exonuclease, very short patch repair endonucleases) (24), transposases (*e.g.,* Tn7 transposase) (25), DNA binding (*e.g.,* RPB5) (26), repair (*e.g.,* MutH) (27), and recombination (*e.g.,* SOX) proteins (28). Based on their genomic location and the flanking genes, Cas4 proteins are of three types: CAS-Cas4 (CRISPR-associated Cas4 genes that occur as part of an array of Cas genes), solo-Cas4 (Cas4 genes located outside CRISPR-Cas locus), and MGE-Cas4 (Cas4 associated with mobile genetic elements) (29). CAS-Cas4s, being associated with CRISPR genes, participate in CRISPR-related adaptive immunity (29). The solo-Cas4s are expected to carry out non-CRISPR functions such as DNA repair and recombination (29), although some studies have shown their involvement in CRISPR function too (21). Likewise, MGE-Cas4 is predicted to be involved in the transposition of mobile genetic elements (29).

Three archaeal Cas4 proteins (SSO0001 and SSO1391 from *Sulfolobus solfataricus* and Pcal_0546 from *Pyrobaculum calidifontis*) have been biochemically characterized Interestingly, these characterized enzymes showed varied oligomeric states: monomer (Pcal_0546), dimer (SSO1391), and decamer (SSO0001) (30–32). Mutational studies revealed that the four conserved cysteine residues are important for Fe-S cluster binding (30, 31). Cas4 are sequence-dependent nucleases, recognizing PAM sequence (17, 23, 25, 33). Although these three characterized proteins were hypothesized to produce ssDNA overhangs, thereby facilitating the insertion of spacers into the CRISPR array, none of the three characterized proteins, SSO0001, SSO1391, and Pcal_0546 have shown a preference for any PAM sequence in the *in vitro* experiments (30–32).

In various CRISPR systems, Cas4 proteins have been reported to coordinate differently in spacer processing. In the type I-A system from *Pyrococcus furiosus*, CAS-Cas4 processes the PAM end while solo Cas4 handles the non-PAM end (21). Similarly, in the type I-A system from *S. islandicus*, Cas4, and CsaI proteins process both ends (34). In the type I-C system from *A. halodurans*, Cas4 processes the PAM end, with the non-PAM end being processed by a cellular exonuclease or Cas1 (35). In general, Cas4, in conjunction with Cas1 and Cas2, is crucial for spacer generation and integration across type I-A, I-B, and I-C subtypes. In the type I-G and type V-B system, a Cas1-4 fusion protein exists, with the Cas4 domain recognizing PAM (36, 37). In summary, the Cas4 protein appears to show PAM specificity only with Cas1 and Cas2 (35, 37, 38).

While several Cas4 proteins have been studied *in vitro* and *in vivo*, their function remains elusive due to significant functional variability. Notably, even among the CAS-Cas4s, their role varies according to different CRISPR subtypes. Incidentally, all of the three *in vitro* characterized archaeal Cas4 proteins (SSO0001, SSO1391, Pcal_0546) were later shown to belong to the solo class of Cas4. Remarkably, no CAS-Cas4 protein of archaeal origin has been characterized *in vitro* until now. Recently, a bacterial Cas4-Cas1 fusion protein (GsCas4, Acc. No. Q74H36.1) from G*eobacter sulfurreducens* (of type IG) has been characterized (36). In this context, we present the novel structure-selective endonuclease activity of archaeal CAS-Cas4 protein from *Thermococcus onnurineus*, TON_0321 (protein Acc. No. TON_0321), associated with type IV-C CRISPR cassette.

## Results

### Types of Cas4 systems

Cas4 proteins, members of the PD-(D/E)XK nuclease superfamily, comprise of a RecB domain, three C-terminal cysteines, one N-terminal cysteine, and an Iron-sulfur cluster. We performed a multiple sequence alignment of Cas4 proteins from various archaeal species using MUSCLE (MUltiple Sequence Comparison by Log-Expectation) (41) (Fig. S1A) and generated a phylogenetic tree using the MEGA (Molecular Evolutionary Genetics Analysis) software suite (Fig. S2) (42). A multiple sequence alignment of archaeal Cas4 proteins shows the conservation of amino acid residues characteristic of the PD-(D/E)XK superfamily, RecB motif, and QhXXY domain (Fig. S1A). The domain arrangement observed with archaeal Cas4 proteins is typical of the AddB family of exonucleases, known for their role in bacterial DNA recombination (43). The four Cysteine residues chelating a Fe-S cluster and metal ion-chelating residues at the active site are also conserved across the Cas4 members (Fig. 1B, Fig. S1A).

Next, we examined each protein’s genomic location to identify its flanking genes (Fig. S3). Phylogenetic analysis (42, 44) and genomic location categorized the Cas4 proteins into two distinct clades: the CAS-Cas4 and the solo Cas4 proteins (Fig. S2). The CAS-Cas4 proteins are part of a larger CRISPR system, surrounded by other Cas proteins. For example, in the CRISPR cassette of the protein TON_0321 (Fig. 1C), Cas4 is located next to several other Cas proteins, including Cas6, Cas10, Cas11, Cas7, Cas5, and Cas2. Because of its association with these other proteins, Cas4 in this case is called a CRISPR-associated Cas4 (CAS-Cas4) protein. On the other hand, solo Cas4 proteins are not part of the CRISPR cassette and are not flanked by other Cas proteins. For instance, proteins like Sso0001 and Pcal_0546 are surrounded by genes that aren’t related to Cas proteins (Fig. 1C). Based on their phylogenetic separation into distinct clades, we anticipate that Cas-Cas4 and Solo-Cas4 proteins exhibit functional divergence. The diversity of Cas4 proteins across different systems emphasizes the need to study each variant individually to fully comprehend their distinct roles, biochemical properties, and potential applications. Therefore, we decided to biochemically characterize the CAS-Cas4 protein to better understand how it might differ from the solo types. This study is the first to investigate a CAS-Cas4 protein in depth, giving us a new basis for comparing it to the solo-type Cas4 proteins.

### TON_0321: Cas4 protein from *Thermococcus onnurineus*

The gene for Cas4 protein in *Thermococcus onnurineus* (TON_0321 protein) is present adjacent to a type IV-C CRISPR Cassette, along with the presence of type III effectors (*Cas2*, *Cas5*, *Cas7*, *Cas10*, *Cas11*) (Fig. 1C). Cas4 proteins are often located adjacent to *Cas1* and *Cas2* genes in most CRIPSR systems, together forming the adaptation module (33, 45). Sometimes, *Cas4* is fused with *Cas1*, as seen in subtypes V-B and the type I-G system of *Methanosarcina barkeri* (15, 36, 37, 46–51) (Fig. 1A)*. Cas4* and *Cas2* (but not *Cas1*) are present next to the type IV-C CRISPR Cassette of *T. onnurineus* (Fig. 1C). Since Cas1, Cas2, and Cas4 form a functional module in many CRISPR subtypes, and Cas1 is absent altogether in the *T. onnurineus* type IV-C CRISPR cassette, we examined the potential interaction between TON_0321 and Cas2 protein from *T. onnurineus*. Interestingly, we did not observe a stable interaction between TON_0321 and Cas2 protein under *in vitro* conditions (Fig. S4). This intriguing outcome led us to explore whether TON_0321 in *T. onnurineus* is a putative protein or exhibits a previously unexplored biochemical profile.

We assessed the integrity of the catalytic site in TON_0321 by predicting its secondary structure, identifying disordered regions (Fig. S5), and modeling its tertiary structure using AlphaFold2 (52, 53). The predicted structure of TON_0321 was then compared with those of related proteins, including Pcal_0546 from *Pyrobaculum calidifontis* (PDB: 4R5Q), SSO0001 from *Sulfolobus solfataricus* (PDB: 4IC1), and GsCas4 from *Geobacter sulfurreducens* (PDB: 7MI4) (Fig. S6). Notably, Cas4 proteins, whether solo or CRISPR-associated, exhibit a similar fold, sharing a conserved core motif with slight variations in the loop regions (Fig. S7). Furthermore, a structural comparison of the catalytic center of TON_0321 with λ exonuclease (PDB: 1AVQ), a well-studied representative member of the PD-(D/E)XK nuclease superfamily, established that the catalytic center of TON_0321 exhibits the typical signature geometry characteristic of PD-(D/E)-XK nucleases (Fig. 1D). In the PD-(D/E)XK nuclease superfamily, to which λ exonuclease belongs, two metal ions (Mg^2+^) in the active site are crucial for catalysis: one activates a water molecule to attack the scissile phosphate while the other stabilizes the transition state (54, 55). The divalent metal ions are chelated by conserved histidine, aspartate and glutamate residues of the RecB motif. In TON_0321, D96 and E105 are identified as active site residues through sequence alignment (Fig. S1A) and conservation of the active site (Fig. 1D). The two residues were mutated to Alanine (TON_0321^D96A/E105A^) and used as a catalytically inactive enzyme in subsequent experiments as a negative control (Fig. S1B). In summary, although TON_0321 does not form a stable complex with Cas2 under the *in-vitro* conditions explored in the current study, it retains the signature residues associated with Cas4 proteins and adopts a fold consistent with other Cas4 family members. The absence of a detectable interaction between TON_0321 and Cas2 is unlikely to result from common artifacts, such as tag interference (Fig. S4E). However, we cannot exclude the possibility that a Cas2–Cas4 interaction may occur in the presence of additional factors under in vivo conditions. Taken together, these findings suggest that TON_0321 from *T. onnurineus* may be a catalytically active nuclease, prompting further investigation into its nuclease activities.

### Optimum catalytic conditions for TON_0321

The TON_0321 protein was purified using affinity chromatography followed by size-exclusion chromatography. To comprehensively characterize the nuclease potential of TON_0321, we optimized a wide range of conditions encompassing protein concentrations, temperature, pH, salt concentrations, and metal ions using single-stranded DNA labeled with Cy5 at the 3′-end. TON_0321 proved to be thermostable and catalytically active over a wide range of temperatures, pH, and NaCl concentrations. The high thermostability of TON_0321 reflects the thermophilic characteristic of *T. onnurineus*. Cas4 enzymes typically follow a divalent metal ion-based nucleophilic attack mechanism to break a phosphodiester bond (54, 56, 57). Nuclease assays were then conducted at either 35 ℃, to preserve the secondary structure of DNA substrates, or 55 ℃, to ensure a fully extended state of single-stranded DNA. A pH of 8.0 and 125 mM NaCl concentration were selected for subsequent experiments based on the stability of protein stability and the annealed synthetic DNA substrates. Additionally, a concentration of 2.5 mM MgCl_2_ was used for the nuclease assays.

### TON_0321 exonuclease activity

We explored the exonuclease activity of TON_0321 using two ssDNA substrates labeled with fluorophores: one labeled with 6-FAM at the 5′ end (Y0-1) and the other with Cy5 at the 3′ end (Y0-4). Experiments were conducted at 35 °C and 55 °C. For the 5′ 6-FAM labeled DNA, Y0-1, we observed a major product of 5-6 bases within 2.5 minutes, increasing over time, indicating 5′ to 3′ exonuclease activity. Similarly, for the 3′ Cy5 labeled substrate (Y0-4), we observed a series of products, gradually catalyzing to a smaller fragment of around four bases over time, further supporting 5′ to 3′ exonuclease activity (Fig. 2A, Fig. S8). Using the Y0-1 substrate, intermediate-sized products were observed, which may be attributed to endonuclease activity, as explored in subsequent sections

**Figure 2:**
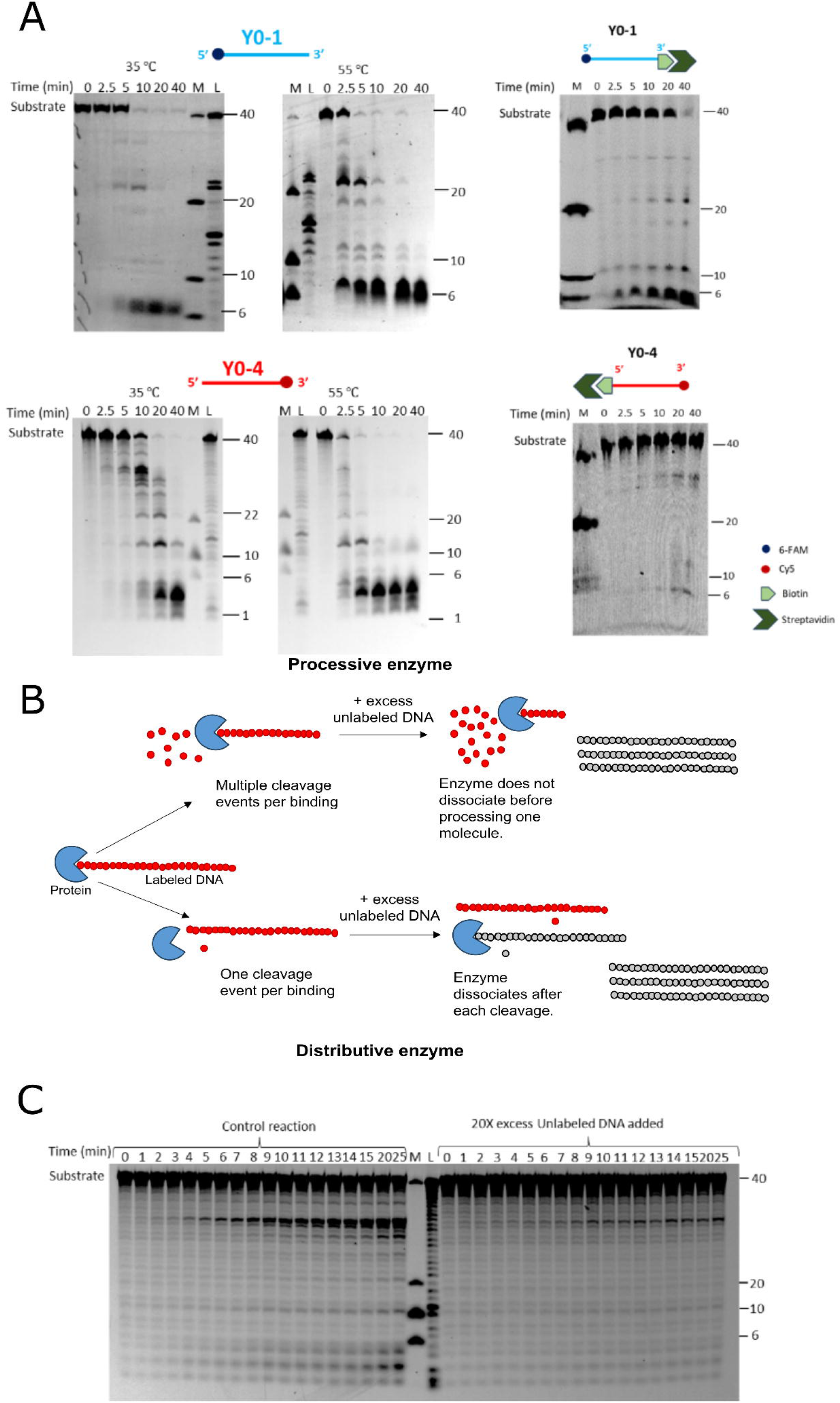
Exonuclease activity of TON_0321 on single-stranded DNA (ssDNA). **(A)** The catalytic activity of TON_0321 on ssDNA labeled at 5′-end with 6-FAM (upper panel) (Y0-1) and ssDNA labeled with Cy5 at 3′-end (lower panel) (Y0-2). The reaction products were resolved on an 18 % TBE-Urea PAGE. Upper panel gels were scanned for 6-FAM signal and the lower panel gels were scanned for Cy5 signal. The activity of TON_0321 on ssDNA labeled at 5′ end with 6-FAM, with biotin at 3′ end (Y0-1_biotin), and ssDNA labeled with Cy5 at 3′ end, and with biotin at 5′ end (Y0-4_biotin). The biotinylated oligos were incubated with 2.5 times molar excess of streptavidin to completely block the biotinylated end of the oligo. The biotin-streptavidin conjugated oligos were used as substrates for activity assay. The reaction products were resolved on an 18 % TBE-Urea PAGE. Gels were scanned for 6-FAM and Cy5 signals. M represents a marker made from mixing synthetic oligonucleotides of different sizes (40, 20, 10 and 6 nucleotides) and L represents a ladder made from DNase digestion of the 40 mer substrate. All experiments were done in triplicates (Fig. S8, Fig. S9). For experiments with ssDNA labeled with Cy5 at 3′end (lower panel), samples for 35℃ and 55℃ were run on the same gel with a common ladder and marker. **(B)** A schematic illustrating the processing of DNA substrate by processive and distributive enzymes. Processive enzymes perform multiple cleavage cycles on a single-bound substrate before releasing it, preventing its binding to a new, unlabeled substrate introduced mid-reaction. Distributive enzymes catalyze a single cleavage per substrate binding, detaching afterward and becoming available to interact with subsequently added substrate molecules. **(C)** TON_0321 acts in a distributive manner. An activity assay reaction was set with 3′ Cy5 labeled ssDNA and formation of products was observed over different time points. In a parallel reaction, 20-fold excess unlabeled DNA of the same sequence was added after two minutes of time point. A decrease in the cleavage of the labeled product in comparison to the control reaction indicated that TON_0321 can dissociate from its substrate and catalyze the reaction in a distributive manner. The reaction products were resolved on an 18 % TBE-Urea PAGE. The gels were scanned for Cy5 signal. M represents a marker made from mixing synthetic oligonucleotides of different sizes (40, 20, 10 and 6 nucleotides) and L represents a ladder made from DNase digestion of the 40 mer substrate. All experiments were done in triplicates (Fig. S11).

To rule out any possibility of 3′ to 5′ exonuclease activity, we used ssDNA substrates blocked with a biotin-streptavidin conjugate at one end and a fluorophore at the other. The 5′ 6-FAM labeled substrate (Y0-1_biotin) was blocked with biotin at the 3′ end, while the 3′ Cy5 labeled substrate (Y0-4_biotin) was blocked with biotin at the 5′ end (Fig. 2A, Fig. S9). These substrates were incubated with streptavidin to form biotin-streptavidin conjugates, followed by an activity assay with TON_0321 at 35 °C. The reactions were analyzed using 18% TBE-Urea PAGE. For Y0-1_biotin, a small (∼6 bp) product accumulated early and with no product formation with Y0-4_biotin, confirming 5′ to 3′ exonuclease activity (Fig. 2A, Fig. S9). A double mutant of metal chelating residues (*i.e.*, D96 and E105), TON_0321^D96A/E105A^ was used as a negative control (Fig. S10).

Next, we aimed to determine how TON_0321 interacts with DNA molecules, specifically whether it acts processively (sliding along the DNA and catalyzing multiple consecutive reactions without releasing the substrate) or distributively (performing only a single reaction with the substrate before releasing it) (58). An exonuclease assay was conducted with ssDNA Y0-4 labeled with Cy5 at the 3′-end at 35 °C. After 1 minute, a 20-fold excess of unlabeled ssDNA was added to the reaction, which was followed for 25 minutes (Fig. 2B). As a control, a similar reaction was performed without adding the excess unlabeled DNA. In this experiment, TON_0321 would have bound the labeled substrate and started its catalysis before adding the unlabeled substrate to the reaction. If the enzyme is processive, the addition of unlabeled substrate would not affect the reaction, and the enzyme would continue to catalyze the labeled substrate it has already bound. Such an enzyme would slide over the bound substrate, completing its catalysis before binding to another substrate molecule. On the other hand, the distributive enzyme would dissociate from the bound labeled substrate molecule after catalyzing one cleavage event. It would have to bind again for the next round of catalysis. In such a case, after the addition of an unlabeled substrate, the enzyme might bind to the unlabeled substrate as it is present in excess, and this would cause a decrease in cleavage of the labeled substrate compared to the control reaction. Interestingly, the addition of unlabeled DNA reduced the cleavage of labeled DNA compared to the control reaction (Fig. 2C, and Fig. S11), indicating that TON_0321 exhibits a distributive mode of action, cleaving once before dissociating from the substrate. In summary, TON_0321 functions as a 5′ to 3′ exonuclease, operating distributively on DNA substrates.

### Endonuclease activity of TON_0321 on dsDNA plasmid

We tested the activity of TON_0321 on linear blunt end dsDNA at 35 ℃ and observed no cleavage, suggesting its inability to cleave dsDNA. However, at 55 ℃, the enzyme showed some catalytic activity, as evident from the accumulation of two fragments sized around 5 bases (major band) and 34 bases (minor band) (Fig. 3A and Fig. S12, Fig. S13). At 55 ℃, the blunt ends of dsDNA could have melted and opened up, creating branches with single-stranded DNA. This branching may have triggered the observed bands, resulting from either exonuclease or endonuclease activity on the branched DNA substrate.

**Figure 3:**
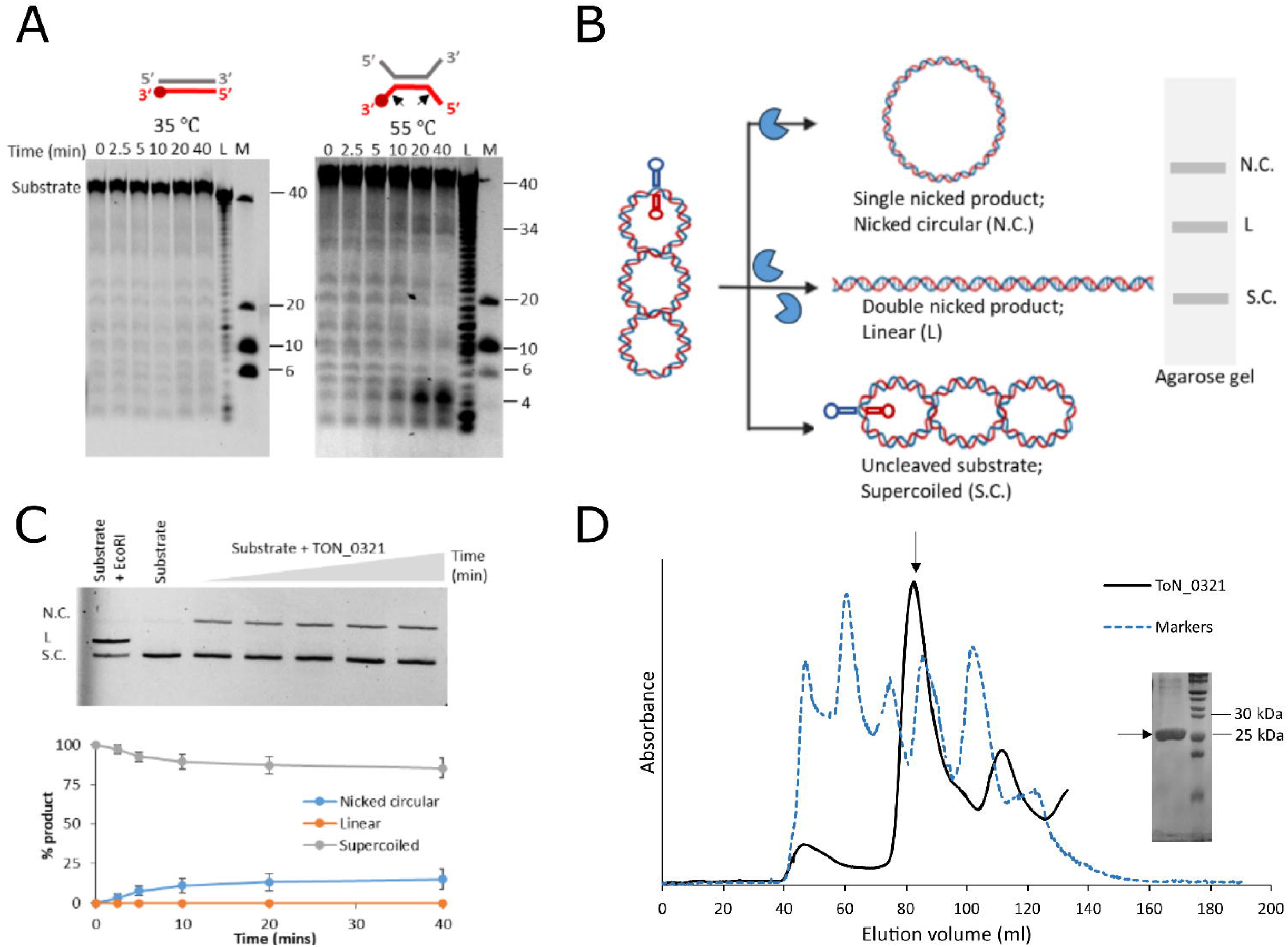
Endonuclease activity of TON_0321. **(A)** Catalytic activity of TON_0321 protein on dsDNA with one strand labeled with Cy5 at the 3′-end at two different temperatures, 35 °C and 55 °C. The reaction products were resolved on an 18 % TBE-Urea PAGE. The gels were scanned for Cy5 signal. M represents a marker made from mixing synthetic oligonucleotides of different sizes (40, 20 10 and 6 nucleotides) and L represents a ladder made from DNase digestion of the 40 mer substrate. All experiments were done in triplicates (Fig. S12). The major cleavage site in the schematic of the DNA substrate is marked by a solid arrow. **(B)** A schematic diagram showing the cruciform assay using the double-stranded plasmid pIRbke8^mut^ containing an inverted repeat sequence that forms a cruciform-type extrusion. The cruciform-like structure on the plasmid contains two sites for EcoRI restriction endonuclease (marked by asterisk). The purified plasmid is in a supercoiled (S.C.) state, which on single nick forms nicked circular (N.C.) DNA and two simultaneous nicks produce a linear (L) form of DNA. **(C)** Ethidium Bromide stained 0.8% agarose gel showing results of cruciform assay carried out with TON_0321 protein. The lane with EcoRI represents positive control and with substrate alone represents negative control. All experiments were done in triplicates (Fig. S14). The graph in the lower panel shows the quantitation of different forms of DNA formed in the cruciform assay along with standard errors calculated from three independent experiments. The reaction was carried out for 40 minutes. Aliquots were taken out at different time points (0, 2.5, 5, 10, 20 and 40 minutes). **(D)** Size exclusion chromatogram of TON_0321 (black) purified on a HiLoad 16/600 Superdex 200 pg gel filtration column along with the gel filtration markers (broken blue) (Vitamin B12, Myoglobin, Ovalbumin, Gamma Globulin, and Thyroglobulin). The peak corresponding to the TON_0321 protein is marked by an arrow. The fractions from this peak were run on a 12% SDS gel. The molecular weight of TON_0321 protein was calculated using a standard curve of markers (Fig. S17C).

To check secondary-structure-guided endonuclease activity and to rule out any contribution from the exonuclease activity of TON_0321, we used a DNA substrate without free ends but featuring secondary structure elements. For this, a cruciform assay was performed with a cruciform containing plasmid pIRbke8^mut^ (substrate 1) (39, 40). The cruciform substrate (pIRbke^8mut^) is a supercoiled plasmid with secondary structure elements, containing an inverted repeat sequence that extrudes into a cruciform structure at 37 °C. A single nick by the nuclease dissolves this structure, producing a nicked circular plasmid, while two nicks result in linear DNA. These forms—supercoiled, nicked circular, and linear—are easily distinguished by agarose gel electrophoresis (Fig. 3B). EcoR1, which linearizes the plasmid; T7 endonuclease 1, which generates both linear and nicked circular plasmids (59); and Nt.BspQ1 (60), a nickase that produces a nicked circular plasmid were used as positive controls (Fig. 3, Fig. S14-S16). Furthermore, a double-stranded DNA plasmid derived from pIRbke8^mut^ but lacking the cruciform structure (substrate 2) was used as a negative control (Fig. S15).

In the cruciform assay, the activity of TON_0321 on the cruciform-containing plasmid (pIRbke8^mut^, substrate 1) resulted in the formation of nicked circular DNA, demonstrating its endonuclease activity on double-stranded DNA. However, negligible product was observed due to the activity of TON_0321 on double-stranded DNA plasmid derived from pIRbke8^mut^ but lacking the cruciform structure (substrate 2) in comparison to substrate 1 (Fig. S15). This indicated TON_0321 requires a DNA branching point to exhibit its endonuclease activity. Since TON_0321 generated a nicked circular DNA as product arising from a single nick, we anticipate it must be a monomer in the solution. In contrast, for linearization of the plasmid two simultaneous and coordinated nicks are essential. Therefore, we determined the oligomeric state of TON_0321 by size exclusion chromatography and found it to exist as a monomer in solution (Fig. 3D and Fig. S17) similar to SSO1391 (31), but unlike Pcal_0546, which is a dimer (31), and SSO0001, which forms a toroidal structure of five dimers (30). This oligomeric state remains unchanged under different protein concentrations (Fig. S17A).

Therefore, TON_0321 can identify branched DNA molecules, exhibit endonuclease activity, and operate as a monomer. In order to further validate the endonuclease activity of TON_0321, we carried out endonuclease assays on various synthetic branched DNA substrates.

### Endonuclease activity of TON_0321 on branched DNA substrates

TON_0321 stands out for its endonuclease activity, a trait not thoroughly explored in other Cas4 proteins. While many PD-(D/E)XK superfamily members exhibit endonuclease capabilities, the Cas4 proteins previously studied (SSO0001, SSO1391, and Pcal_0546) demonstrated endonuclease activity solely on circular ssDNA plasmids, unable to cleave circular dsDNA plasmids (30–32). Prompted by the endonuclease activity of TON_0321 on cruciform plasmids and dsDNA at higher temperatures, we expanded our investigation to include its activity on various branched DNA structures, such as 5′ flap, 3′ flap and splayed arm at 35 ℃ (Fig. 4, Fig. S18, Table S2). TON_0321 efficiently processed all of these branched DNA substrates. In the case of the 5′ flap, TON_0321 cleaved the Cy5 labeled strand two nucleotides from the branching point in the 5′ direction, with no nick in the 6-FAM labeled strand (Fig. 4A, Fig. 4B and Fig. S19). With the 3′ flap substrate, TON_0321 cleaved only the Cy5 labeled strand two nucleotides away from the junction in the 5′ direction (Fig. 4A, Fig. 4B and Fig. S19). In the case of the splayed arm substrate, TON_0321 generated a major nick at two nucleotides away from the branching point in the Cy5 labeled strand, while a minor nick is also observed on the 6-FAM labeled strand as well but during later time points (Fig. 4A, Fig. 4B and Fig. S19). Notably, TON_0321 did not make any major nicks on the 6-FAM labeled strand across all substrates studied (Fig. 4A, Fig. 4B, and Fig. S19). Catalytically inactive TON_0321^D96A/E105A^ was used as a negative control (Fig. S20).

**Figure 4:**
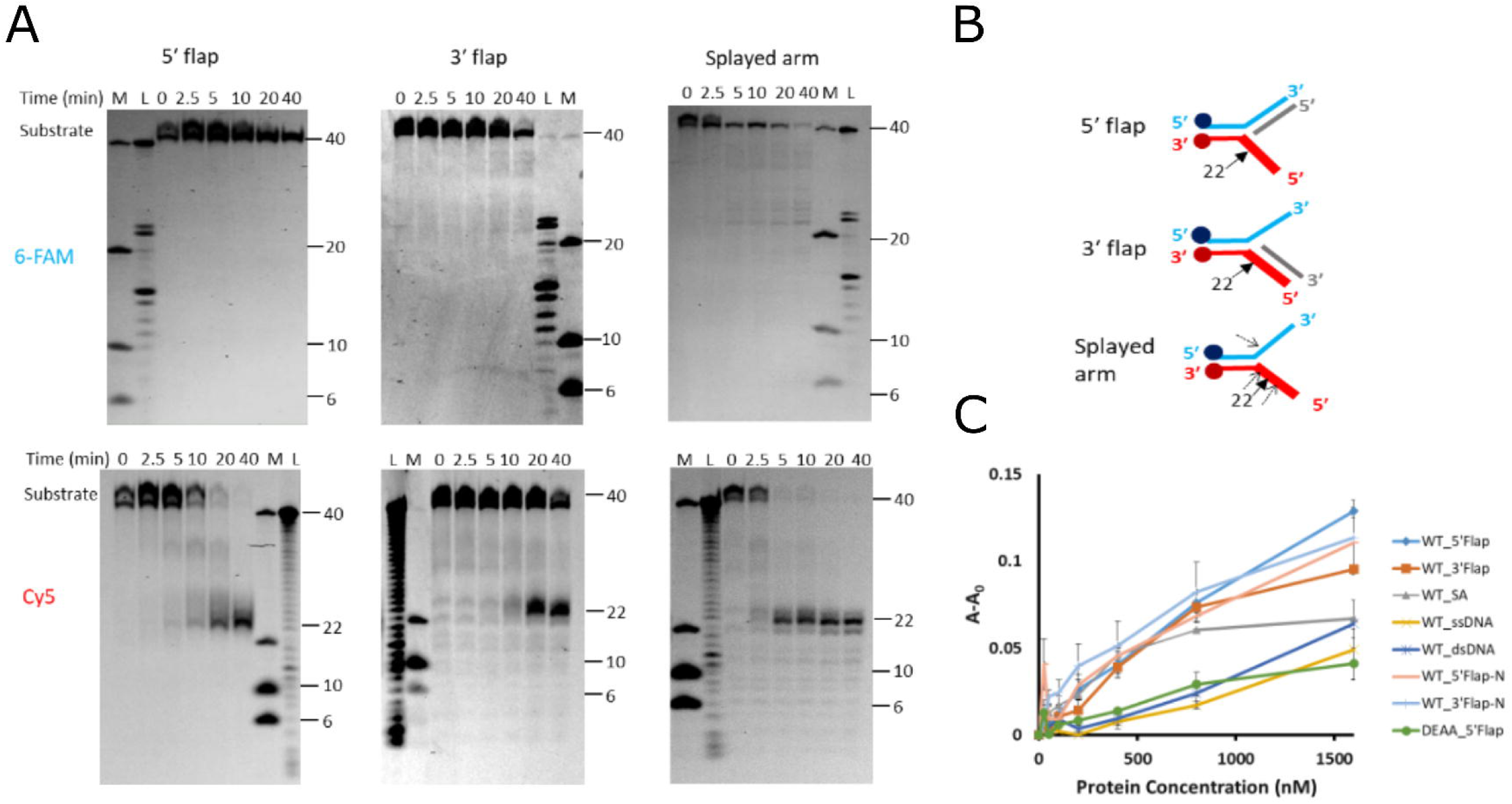
Endonuclease activity of TON_0321 on branched DNA substrates. **(A)** The catalytic activity of TON_0321 protein on branched DNA molecules: 5′ flap, 3′ flap, and splayed arm. Each substrate has two labels: one strand labeled at 5′-end with 6-FAM and another strand labeled at 3′-end with Cy5. The reaction products were resolved on an 18 % TBE-Urea PAGE. The same gel was scanned for 6-FAM signal (upper panel) and for the Cy5 signal (lower panel). M represents a marker made from mixing synthetic oligonucleotides of different sizes (40, 20 10 and 6 nucleotides) and L represents a ladder made from DNase digestion of the 40 mer substrate. All experiments were done in triplicates (Fig. S19). **(B)** The schematic of various branched DNA substrates shows the major cleavage sites as solid arrows and the minor sites as broken arrows. **(C)** Binding study of TON_0321 with different DNA substrates: ssDNA (light blue), dsDNA (orange), 5′ flap (grey), 3′ flap (yellow), and splayed arm (dark blue) using fluorescence anisotropy. DEAA_5’ flap represents binding analysis of catalytically dead TON_0321^D96A/E105A^ in the presence of a 5’ flap substrate. 6-FAM-labeled substrates were used for anisotropy experiments. The Y-axis shows a change in anisotropy (A – A_0_), where A is observed anisotropy and A_0_ is anisotropy of DNA substrate alone.

We used fluorescence anisotropy to assess the binding of TON_0321 with various DNA substrates, both branched and non-branched (Fig. S21A, Table S3). The substrates tested included ssDNA, blunt-end dsDNA, 5′ flap, 3′ flap, and a splayed arm, each labeled with a 5′ 6-FAM label on one of the arms. TON_0321 showed binding with all of these substrates (Fig. 4C, Table S4). Despite no catalytic activity on dsDNA, TON_0321 still exhibited comparable binding to other DNA substrates (Fig. 4C). We also looked into binding of TON_0321 ^D96A/E105A^ with 5’ flap substrate. Notably, this active-site mutant, lacking key metal-chelating residues, exhibited reduced binding compared to the wild-type protein (Fig. 4C and Fig. S21B). In summary, TON_0321 is active with all tested branched substrates (5′ flap, 3′ flap, splayed arm), cleaving only the Cy5 labeled strands. TON_0321 can bind to all DNA substrates, including dsDNA.

### Sequence *vs* structure dependency

The consistent cleavage of the Cy5 labeled strand in all substrates by TON_0321 suggests that this activity might be sequence-dependent or the enzyme recognizes branch points in an orientation-specific manner. Cas4 proteins are known for identifying PAM sequences for spacer acquisition in the adaptation step (20, 21, 35, 37). The type I-E CRISPR system from *E. coli* lacks Cas4 protein and its function is complemented by the C-terminal loop of Cas1 protein as it performs PAM recognition. On the other hand, in other CRISPR types where Cas4 is present, Cas1 protein lacks this C-terminal loop (61). Since the type IV-C CRISPR cassette under study lacks Cas1, we investigated if the Cas4 protein, TON_0321, has a sequence dependency. Given that TON_0321 consistently nicked only the Cy5 labeled strand, we tested this by reversing the sequence near the branch point in new 5′ flap (5′ flap-N) and 3′ flap (3′ flap-N) substrates (Fig. 5 and Fig. S22). Interestingly, TON_0321 continues to nick at the same position, two nucleotides away from the junction point in the 5′ direction, on the Cy5 labeled strand, regardless of the sequence, thus ruling out sequence dependency in the presented experimental set-up.

**Figure 5:**
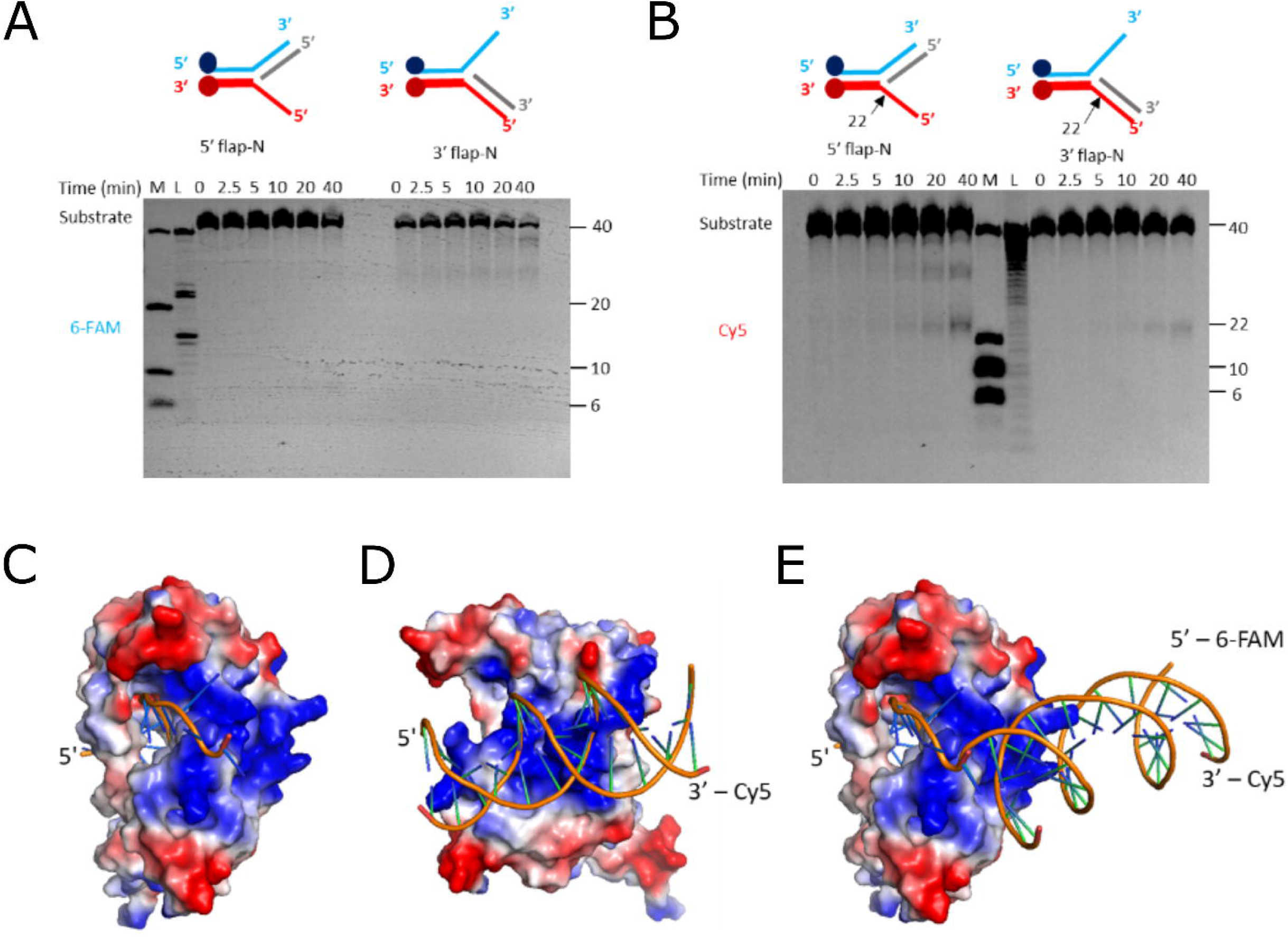
TON_0321 is a secondary structure-selective endonuclease. The catalytic activity of TON_0321 protein on branched DNA molecules, 5′ flap-N and 3′ flap-N, with each having one strand labeled at 5′-end with 6-FAM and another strand labeled at 3′-end with Cy5. The reaction products were resolved on 18 % TBE-Urea PAGE. The gels were scanned for **(A)** 6-FAM signal and **(B)** Cy5 signal. M represents a marker made from mixing synthetic oligonucleotides of different sizes (40, 20, 10, and 6 nucleotides), and L represents a ladder made from DNase digestion of the 40 mer substrate. All experiments were done in triplicates (Fig. S22). The major cleavage site in the schematic of branched DNA substrates is marked by a solid arrow. DNA binding model of TON_0321 with **(C)** ssDNA, **(D)** dsDNA, and **(E)** Splayed arm substrate. The bound DNA was modeled by superimposing the catalytic domain of TON_0321 with the bound structure of GsCas4 (PDB: 7MI4).

Next, we checked if the unique nicking of one specific arm in branched DNA substrates (*i.e*., the Cy5 labeled arm) by TON_0321 relates to its orientation-specific recognition at the branch point. Using AlphaFold2, we modeled the structure of TON_0321, with high-confidence prediction scores (pLDDT value of 87.44) (52, 53). The model showed a positively charged patch positioned at an angle to the catalytic site, capable of binding double-stranded DNA (dsDNA). However, for DNA to reach the catalytic site for cleavage, it must bend, a flexibility more easily achieved at the branched points of DNA (Fig. 5C-5E, Fig. S23). Many structure-selective endonucleases are known to utilize DNA bending and positively charged surface patches to carry out catalysis near the branching point (62–64). We further modeled single-stranded DNA (ssDNA) in the catalytic site of TON_0321 using GsCas4 (PDB: 7MI4) and connected it to the dsDNA bound at the positively charged patch, creating a splayed arm structure (Fig. 5E). This model suggests that DNA bending is crucial for branched DNA recognition and the selective cleavage of the Cy5-labeled arm (Fig. 5E). Cleavage of the 6-FAM-labeled arm would require reverse DNA orientation (impacting the stereochemistry at the catalytic site), or reorientation of the dsDNA and branch point, as seen in PDB: 7MI4. Unlike PDB: 7MI4′s structure, TON_0321 lacks stabilizing elements for dsDNA. Therefore, considering the observed catalytic behavior and our model predictions, it becomes clear that TON_0321 selectively targets branched DNA molecules.

## Discussion

CAS-Cas4 proteins have been established to participate in the adaptation step, possibly through recognition of PAM sequences, thereby impacting the correct integration of functional spacers. However, their mode of action and specific role appear to be quite varied between various CRISPR systems/types. This is not surprising, given the high sequence variability among the Cas4 proteins. The phylogenetic studies have suggested that CAS-Cas4 proteins do exhibit CRISPR-type specific congruence in their evolutionary path (26), suggesting that the Cas4 of a particular CRISPR type/subtype should exhibit unique and specific sequence and structural features. Characterization of CAS-Cas4 proteins from different CRISPR types would aid in correlating their sequence and structural features with their experimentally observed function divergence. To date, three Cas4 proteins have been characterized biochemically from the archaeal systems: SSO0001 and SSO1391 from *S. solfataricus* and Pcal_0546 from *P. calidifontis*. Unlike TON_0321, none of the three Cas4 proteins belong to any CRISPR type but belong to the solo class of Cas4 proteins (30–32) (Fig. 1). Only a Cas4-Cas1 fusion protein from the bacteria *Geobacter sulfurreducens,* Q74H36.1, belonging to CRISPR type I-G, has been recently characterized (36). Although Cas4 proteins are typically absent in type IV CRISPR systems (65, 66), the presence of a Cas4 gene (TON_0321 protein) adjacent to a type IV-C CRISPR system in *T. onnurineus* marked an intriguing deviation from this pattern. Furthermore, the presence of *Cas2* and the absence of *Cas1* genes, and the inability of TON_0321 to form a stable complex with Cas2 protein further underscores the intricate diversity and complexity inherent in Cas4 proteins across various organisms and CRISPR systems and warrants biochemical characterization.

Our studies have established that, as a PD-(D/E)XK superfamily member, the TON_0321 protein demonstrates characteristic divalent metal ion-dependent nuclease activity. TON_0321 exhibits 5′ to 3′ exonuclease activity akin to SSO0001 and Pcal_0546, while SSO1391 possesses both 5′ to 3′ and 3′ to 5′ exonuclease activities. In the present study, TON_0321 is classified as an exonuclease based on its requirement for a free 5’ end, and the property of cleaving the substrate in a stepwise manner, even though it does not produce mononucleotides or dinucleotides as products, but produces oligonucleotides instead. This feature, *i.e.*, generation of oligonucleotides, is notably distinct from other archaeal Cas4 proteins characterized so far. TON_0321 does not degrade circular DNA unless it contains a branched structure. In this respect, TON_0321 is similar to other known exonucleases such as *E. coli* Exonuclease V (RecBCD), Exonuclease VII, and phage T5 exonuclease, all of which generate short oligonucleotide products while acting from DNA termini (67–70).

Exploring the endonuclease potential of TON_0321 offers further valuable insights into its unique characteristics. TON_0321 did not cleave linear dsDNA at a lower temperature (35 °C) but it could cleave the dsDNA at a higher temperature (55 °C) due to the possible opening of the blunt ends. It also cut circular dsDNA with secondary structures, suggesting a new ability to recognize and cut branched DNA substrates, not seen in other archaeal Cas4 proteins (30–32) (summarized in Table 1). Additionally, TON_0321 can cut several branched DNA molecules like 5′ flap, 3′ flap, and splayed arm, marking the first instance of a Cas4 protein being identified as a structure-dependent endonuclease. Most of the structure-dependent endonucleases possess structural features to identify branching in DNA substrates. Many of them recognize bending in DNA. The predicted structure of TON_0321 shows a highly positively charged surface at a sharp angle with the catalytic site allowing the possibility of recognizing sharp bends in branched DNA substrates. This is also reflected in the observation where TON_0321 can bind a dsDNA without catalyzing it but at the same time can catalyze many branched DNA substrates.

None of the previously characterized Cas4 proteins have shown sequence-dependent cleavage *in vitro* (30–32). On similar lines, TON_0321 also showed sequence non-specific endonuclease activity in the present context. Cas4 proteins from type I-A, I-B, and I-C systems, without other adaptation proteins (Cas1 and Cas2), also show sequence non-specific nuclease activity *in vivo* (35, 38) suggesting that the interaction with Cas1 and Cas2 may be essential for PAM recognition-based activity (38). The type IV-C CRISPR Cassette under study lacked Cas1 protein from the adaptation module. However, we cannot rule out the possibility of TON_0321 forming a complex with adaptation protein Cas2 *in vivo*, activating its sequence specificity, though under *in vitro* conditions, we did not obtain a stable complex between TON_0321 and Cas2.

Another significant difference between TON_0321 and other characterized Cas4 proteins lies in their oligomeric state. SSO0001 forms a toroidal structure from five dimers (30), SSO1391 exists as a dimer in solution (31), and Pcal_0546 exists as a monomer (31). Interestingly, TON_0321 is also monomeric in solution at several different concentrations tested (Fig. S17). SSO0001, due to its toroidal structure, has been proposed to serve as a sliding clamp for other Cas proteins (30). Superposition with the toroidal structure of SSO0001 suggests that the N-terminus of TON_0321 is oriented away from the protein–protein interaction interface, making steric hindrance from an N-terminal tag during oligomerization unlikely based on the current model. However, this interpretation remains subject to the limitations inherent in structural modelling (Fig. S17B). Also, SSO0001 might serve as a powerful DNA-degrading machinery if its multiple active sites can function in unison (30). Since Cas4 proteins typically function in conjunction with other adaptation proteins, such as Cas1 and Cas2, as observed for monomeric Pcal_0546, it is plausible that TON_0321 also forms a complex with other adaptation proteins. Future studies could investigate whether TON_0321 participates in CRISPR-related processes, such as generating single-stranded DNA overhangs in spacers, or explore its potential roles in non-CRISPR functions, such as DNA repair or broader nucleic acid metabolism (31).

Studying the uniqueness of TON_0321 has enhanced our understanding of the functional divergence of Cas4 proteins, and expanded the landscape of nuclease activity hitherto assigned to Cas4 proteins. TON_0321 is a versatile nuclease enzyme capable of processing single-stranded DNA and branched-chain substrates, suggesting its potential involvement in diverse cellular pathways requiring precise nucleic acid processing. TON_0321 shows endonuclease activity towards characteristic intermediates of DNA repair pathways, chromosome segregation mechanisms, and CRISPR systems. Like TON_0321, Flap endonuclease 1 (FEN1) also exhibits 5′ to 3′ exonuclease and structure-selective endonuclease activities (71, 72). The structure-selective endonuclease activity of TON_0321 highlights its potential as a structure-dependent nuclease, with its ability to target branched DNA intermediates offering opportunities to complement existing CRISPR-Cas technologies and inspire the development of innovative genome editing tools. Furthermore, TON_0321 may play a significant role in plasmid propagation by enhancing recombination with other nucleic acids or maintaining plasmid mobility, warranting further investigation (66).

The unique properties of TON_0321 may also offer valuable insights for developing innovative genome editing tools, expanding the toolkit for genetic engineering and molecular biology applications. Future efforts to decouple its endonuclease and exonuclease functions, through targeted single or combinatorial mutagenesis combined with substrate-specific nuclease assays and stoichiometric binding analysis, could yield deeper mechanistic insights. Understanding this bifunctionality at the molecular level could have broader implications for manipulating Cas4-family enzymes in genome engineering and DNA repair contexts. In summary, adding upon the information available for Cas4 proteins, the current study delves into the characteristics of the TON_0321 Cas4 protein, located next to the Type IV-C CRISPR cassette. Hereby, TON_0321 is a structure-selective endonuclease with 5′ to 3′ ssDNA-specific exonuclease activity.

## Experimental Procedures

### Protein expression, purification and mutagenesis

The genes encoding for TON_0321 protein and Cas2 protein (protein Acc. No.: TON_0320) were amplified from the genomic DNA of *Thermococcus onnurineus NA1* using gene-specific primers (SI Table 1). The gene for TON_0321 was cloned in the pGEM®-T cloning vector (Promega Corporation) and then, subcloned in pET28a vector with N-terminal 6 X His tag (Novagen) and transformed into BL21 competent cells (Thermo Fisher Scientific). The gene for Cas2 was cloned into pGEX-4T vector. For the active site mutant of TON_0321, D96, and E105 amino acids were mutated to Alanine using the Quick-change site-directed mutagenesis via KOD hot start DNA polymerase (Merck Millipore). For TON_0321 expression, transformed bacteria were grown till OD_600_ reached ∼ 0.6 in LB media containing 50 µg ml^-1^ kanamycin at 37 ℃, and then, induced with 1 mM IPTG at 37 ℃ for 4 hours. Bacteria were pelleted at 4000 rpm for 10 minutes at 4 ℃. The expressed protein was purified using Ni-NTA affinity chromatography. The bacterial pellet from the 2-litre culture was resuspended in lysis buffer containing 25 mM tris-HCl (pH 8.0), 10% glycerol, 500 mM NaCl, and 5 mM imidazole supplemented with PMSF and lysozyme. The suspension was incubated at 37 ℃ for 30 minutes, followed by sonication. The cell lysate was centrifuged at 10, 000 rpm for 20 minutes, and the collected supernatant was loaded onto a pre-equilibrated Ni-NTA column (Bio-Rad). After binding (binding buffer: 25 mM Tris-HCl (pH 8.0), 10% glycerol, 500 mM NaCl, and 25 mM imidazole) for 2 hours, the Ni-NTA beads were firstly washed with wash buffer containing 25 mM Tris-HCl (pH 8.0), 10% glycerol, 500 mM NaCl, and 50 mM imidazole and then, eluted in 25 mM Tris-HCl (pH 8.0), 500 mM NaCl, 10% glycerol, and 300 mM imidazole. Fractions obtained were run on a 12% SDS-PAGE gel. The eluted fraction was then, loaded on a HiLoad 16/60 S200 column (GE Healthcare) pre-equilibrated with 25 mM tris-HCl (pH 8.0), 10% glycerol, 500 mM NaCl. The peak obtained was confirmed for the presence of protein by running the fractions on a 12% SDS-PAGE gel. The protein fractions were stored in 50% glycerol at -20 ℃. The active site mutant protein TON_0321^D96A/E105A^ was purified similarly.

For Cas2 expression, transformed bacteria were grown till OD_600_ reached ∼ 0.6 in LB media containing 100 µg ml^-1^ Ampicillin at 37 ℃. Protein expression was induced with 0.4 mM IPTG, followed by overnight incubation at 16 °C. The culture was then centrifuged at 6000 rpm for 20 minutes at 4 °C to harvest the cells. For Cas2 purification, the cell pellets from the 2L culture were resuspended in lysis buffer (25 mM Tris-HCl (pH 8.0), 150 mM NaCl, 10% glycerol) containing lysozyme and PMSF, and incubated on ice for 30 minutes. The suspension was then sonicated and centrifuged at 18, 000 rpm for 30 minutes at 4 °C. The supernatant was loaded onto a GSTrap™ 4B column (Cytiva) pre-equilibrated with wash buffer (25 mM Tris-HCl (pH 8.0), 150 mM NaCl, 10% glycerol). Elution was performed using elution buffer (25 mM Tris-HCl (pH 8.0), 150 mM NaCl, 10% glycerol, 10 mM glutathione) on an ÄKTA FPLC system. Fractions containing the protein of interest were concentrated using a 10 kDa Amicon (MERCK). For further purification, size exclusion chromatography was performed by injecting the concentrated protein into a HiLoad 16/60 S200 column (GE Healthcare) equilibrated with buffer (25 mM Tris-HCl (pH 8.0), 150 mM NaCl, 5% glycerol). The chromatogram peaks were analyzed on a 12% SDS-PAGE, and the fractions containing the pure protein of interest were concentrated and stored at -20°C in 50% glycerol.

### Nuclease assay

The HPLC-purified ssDNA oligonucleotides were purchased from SIGMA (SI Table 2A). The DNA substrates were prepared by annealing these ssDNA oligonucleotides in the combination mentioned in SI Table 2B. For annealing, different ssDNA oligonucleotides were mixed in equimolar ratio, and heated at 95 ℃ for 5 minutes followed by slow cooling overnight. The substrate for the nuclease reaction comprised of a mixture of 100 nM unlabeled substrate and 25 nM of labeled substrate. The labeled substrate consisted of 6-FAM on the 5′-end of Y0-1 and Cy5 on the 3′-end of Y0-4. The reaction mixture without protein, consisting of 50 mM Tris-HCl (pH 8.0), 2.5 mM MgCl_2_, 1.0 mM DTT, 125 mM NaCl, and 0.1 mg/ml BSA, was incubated at 35 ℃ for 15 minutes. To initiate the reaction 25 nM of protein was added. Aliquots were taken out at different time points (0, 2.5, 5, 10, 20, and 40 minutes) and the reaction was quenched using 5 mM EDTA, 2 mg/ml Proteinase K, and 0.2% SDS treatment of reaction aliquots. The samples were resolved on 18% TBE Urea PAGE and scanned using a Typhoon scanner (GE Healthcare). The quantification of bands (substrate and products) was performed using Amersham ImageQuant software, involving lane creation, background subtraction, band detection, and relative intensity quantification. The samples for Urea PAGE were prepared by heating the reaction mix at 95 ℃ for 10 min in formamide dye. Different DNA oligonucleotides were used to generate a DNA ladder (SI Table 2C). All experiments were done in triplicate.

### Cruciform assay

The cruciform assay was performed using plasmid pIRbke8^mut^ (39, 40). The cruciform substrate (pIRbke8^mut^) is a supercoiled DNA plasmid without free ends that contains secondary structure elements. Plasmid pIRbke8^mut^ (substrate 1) includes an inverted repeat sequence, which, upon incubation at 37 °C, forms a cruciform-like structure resembling a four-way junction. This cruciform-like structure was eliminated through site-directed mutagenesis (primers used are mentioned in SI table 1), resulting in a double-stranded, supercoiled plasmid without free ends or secondary structure (substrate 2). The plasmids were transformed into DH5α competent cells and positive transformants were selected on LB plates containing ampicillin. A primary culture was set and harvested when OD_600_ reached 0.5 (mid-log phase). The plasmid was isolated from the culture pellet using a Qiagen® midiprep plasmid isolation kit. For each experiment, both the plasmids were diluted to 10 nM with water. The reaction was performed with 2 nM of plasmid DNA in a buffer containing 50 mM Tris-HCl (pH 8.0), 2.5 mM MgCl_2_, 1.0 mM DTT, 125 mM NaCl and 0.1 mg/ml BSA incubated at 37 ℃ for 30 minutes to induce cruciform extrusion. To initiate the reaction, 25 nM of protein was added, and the reaction was carried out for 40 minutes. Aliquots were taken out at different time points (0, 2.5, 5, 10, 20 and 40 minutes). EcoRI, T7 endonuclease I, and Nt.BspQ1 enzymes from New England Biolabs were used as positive controls, and the reactions were set for 40 minutes. The reactions were quenched using 5 mM EDTA, 2 mg/ml Proteinase K, and 0.2% SDS. Products were run on 0.8% agarose gel. Gel was stained with EtBr (0.5 µg/ml in 1X TBE) followed by destaining using water and visualized using Gel doc XR+ system (Bio-Rad). The quantification of bands (substrate and products) was performed using Image Lab (Bio-Rad) software. All experiments were done in triplicate. Figure 3B created in BioRender. Gaur, V. (2025) https://BioRender.com/sj9qxon and Figure S15A created in BioRender. Gaur, V. (2025) https://BioRender.com/rj3p8sh.

### Fluorescence anisotropy

To study the binding of TON_0321 protein with various DNA substrates (Table S3), fluorescence anisotropy was used. DNA substrates were labeled with 6-FAM on 5′-end of Y0-1. 35 nM of DNA substrate mixture consisting of 10 nM unlabeled DNA and 25 nM of labeled DNA was used. Protein concentration varied in the range of 0 to 1600 nM. For DNA substrate binding studies, since we believe metal ions can also contribute to DNA binding, instead of using Ala mutants, we suppressed catalytic activity by using Zn^2+^ ions in the DNA substrate binding studies. The reaction was set up in a buffer containing 20 mM Tris-HCl (pH8.0), 100 mM NaCl, 0.5 mM DTT, 0.1 mg/ml BSA, and 20 mM ZnSO_4_ at 25 ℃ in 96 well flat bottom black polystyrene plates (BRAND®). Anisotropy was measured using a POLARstar Omega plate reader at an excitation wavelength of 485 nm and an emission wavelength of 520 nm. Fluorescence anisotropy was calculated as (I_||_ – I_⊥_) / (I_||_ + 2I_⊥_), where I_||_ and I_⊥_ are intensities in parallel and perpendicular directions. Binding was studied as the change in anisotropy (A-A_0_) versus protein concentration, where A is the observed anisotropy, and A_0_ is the anisotropy of substrate alone. All experiments were done in triplicate.

### Oligomeric state determination

To determine the oligomeric state of TON_0321, size exclusion chromatography was used. HiLoad 16/60 S200 column (GE Healthcare) pre-equilibrated with 25 mM Tris-HCl (pH 8.0), 10% glycerol, 500 mM NaCl. The column was calibrated using gel filtration markers (Bio-Rad, Cat. No. 151-1901). A standard curve was generated for the gel filtration markers (Thyroglobulin, Vitamin B12, Myoglobin, Ovalbumin, and Gamma Globulin). Kav was calculated as (Ve – Vo) / (Vt – Vo), where Ve, Vo and Vt are elution volume, void volume, and total volume, respectively. The column′s void volume was determined using Thyroglobulin (670 kDa).

### *In vitro* double pulldown assay

75 μg of each TON_0321, and Cas2 were mixed and incubated with 100 μl of glutathione Sepharose 4B beads for 2 hours at 4 °C (washed and preincubated with 25 mM Tris-HCl pH 8.0, 150 mM NaCl, and 5% Glycerol). After incubation, the protein-bead mixture was centrifuged using a Corning® Costar® Spin-X® centrifuge tube filter at 6000 rpm for 2 minutes at 4 °C, and the flow-through was collected. The beads were then washed 10 times with wash buffer (25 mM Tris-HCl pH 8.0, 150 mM NaCl, and 5% Glycerol). Proteins were eluted from the beads using an elution buffer (25 mM Tris-HCl pH 8.0, 150 mM NaCl, 5% Glycerol, and 10 mM Glutathione). The eluted protein was incubated with 100 μl of Ni-NTA agarose beads (QIAGEN) (washed and preincubated with 25 mM Tris-HCl pH 8.0, 150 mM NaCl, and 5% Glycerol) for 2 hours at 4 °C with gentle shaking. Following incubation, the protein-bead mixture was centrifuged again using a Corning® Costar® Spin-X® centrifuge tube filter at 6000 rpm for 2 minutes at 4°C, and the flow-through was collected. The beads were extensively washed 10 times with wash buffer (25 mM Tris-HCl pH 8.0, 150 mM NaCl, and 5% Glycerol). Finally, proteins were eluted by adding elution buffer (25 mM Tris-HCl pH 8.0, 150 mM NaCl, 5% Glycerol, and 500 mM imidazole. The flow-through, wash, and elution fractions were analysed on Bio-Rad 4-20% Mini-PROTEAN® TGX™ Precast Protein gels stained with Coomassie Brilliant Blue.

## Data Availability

All the data described in the manuscript are present in the manuscript or the supporting information.

## Supporting Information

This article contains supporting information (73–76).

## Acknowledgements

We are thankful to the CIF facilities at NIPGR and University of Delhi, South Campus. The authors thank Director, NIPGR for constant support. We are thankful to Dr. Dinakar Salunke and Dr. Deepshika Malik for the critical reading of the manuscript.

## Author contributions

M.J., A.K.P., S.A., P.R., and K.J. investigation; S.C. resources; M.J., V.G., and M.G. formal analysis; M.J. writing - original draft; M.J., V.G., and M.G. writing – review & editing; V.G. and M.G. conceptualization; V.G. and M.G. project administration; V.G. and M.G. funding acquisition.

## Funding

NIPGR core grant (to V.G.); DBT Ramalingaswami Fellowship [BT/RLF/Re-entry/27/2017] (to V.G.); SERB CRG [CRG/2020/000335] (to V.G.); SERB CRG [CRG/2019/001310] (to M.G.); IOE research grant (IOE/2021/12/FRP) (to M.G.) and (IOE/2023-2024/12/FRP) (to M.G.).

## Conflict of Interests

The authors declare that they have no conflict of interest with the contents of this article.

## Abbreviations

6-FAM: (6-carboxyfluorescein)
bp: (base pairs)
Cas: (CRISPR-associated protein)
CAS-Cas4: (CRISPR-associated Cas4 gene)
CRISPR: (Clustered Regularly Interspaced Short Palindromic Repeats)
crRNA: (CRISPR RNA)
Cy5: (Cyanine-5 fluorophore)
dsDNA: (double-stranded DNA)
MEGA: (Molecular Evolutionary Genetics Analysis)
MGE-Cas4: (Mobile Genetic Element–associated Cas4)
MUSCLE: (MUltiple Sequence Comparison by Log-Expectation)
PAM: (Protospacer Adjacent Motif)
PDB: (Protein Data Bank)
PD-(D/E)XK: (Proline–Aspartate–(Aspartate/Glutamate)–X–Lysine endonuclease superfamily)
pLDDT: (Predicted Local Distance Difference Test)
RecB: (Recombination protein B domain)
ssDNA: (single-stranded DNA).

## Notes

### Competing Interest Statement

The authors have declared no competing interest.

### Summary of Updates

Additional data has been created and the revised manuscript has additional data.

