## Supplemental Information for "Branched DNA Processing by a Thermostable CAS-Cas4 from *Thermococcus onnurineus*: Expanding Biochemical Landscape of Nuclease Activity"

*Corresponding authors

**List of the material included**

Figures: Fig. S1 to Fig. S23

Tables: Table S1 to Table S4

A

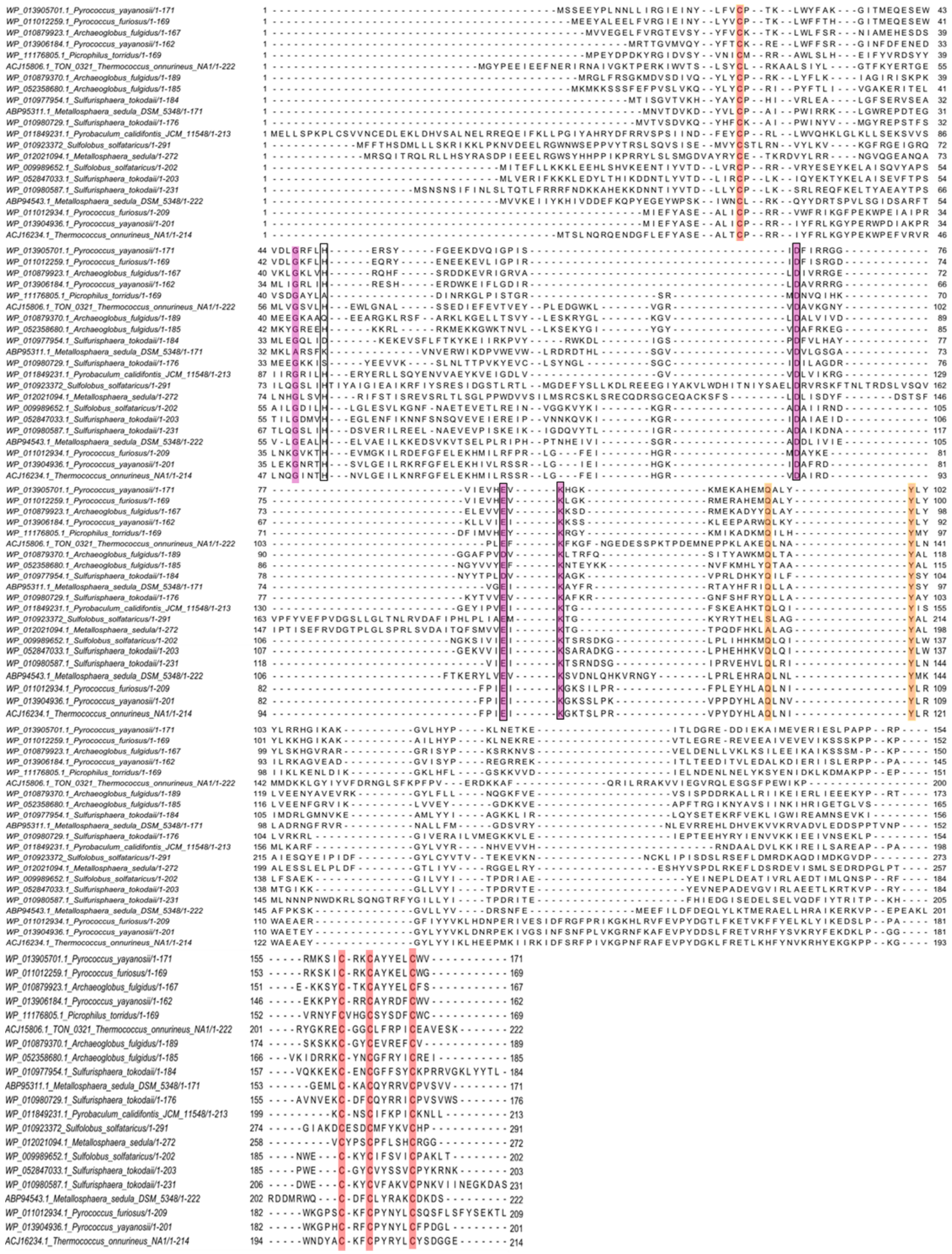

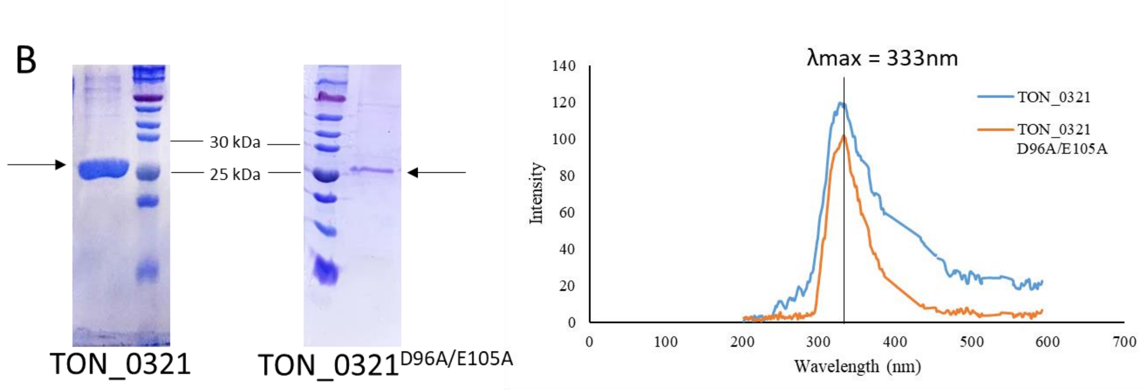


**Figure S1. (A) Multiple sequence alignment.** Multiple sequence alignment of representative members of the archaeal Cas4 protein family using MUSCLE (73) and visualization with Jalview (74) . The NCBI accession numbers of proteins followed by organism names are used for nomenclature of protein sequences. The conserved residues involved in binding Fe-S cluster are highlighted in red color, the RecB motifs I, II, and III are highlighted in magenta color, the QhxxY domain is highlighted in orange color and the metal ion coordinating residues are marked by a black box. (B) Purification profile of TON_0321 wild type and mutant TON_0321 $^{D96A/E105A}$ protein. SDS gel showing the purified TON_0321 wild type protein and the mutant protein TON_0321 $^{D96A/E105A}$. Arrow marks the presence of protein bands. The SDS-PAGE profile of TON_0321 is also used in Fig. S4B and Fig. 3. Fluorescence spectra of the wild type and mutant protein showing the absorption peak at 333 nm for both indicating similar structural fold and conformation.

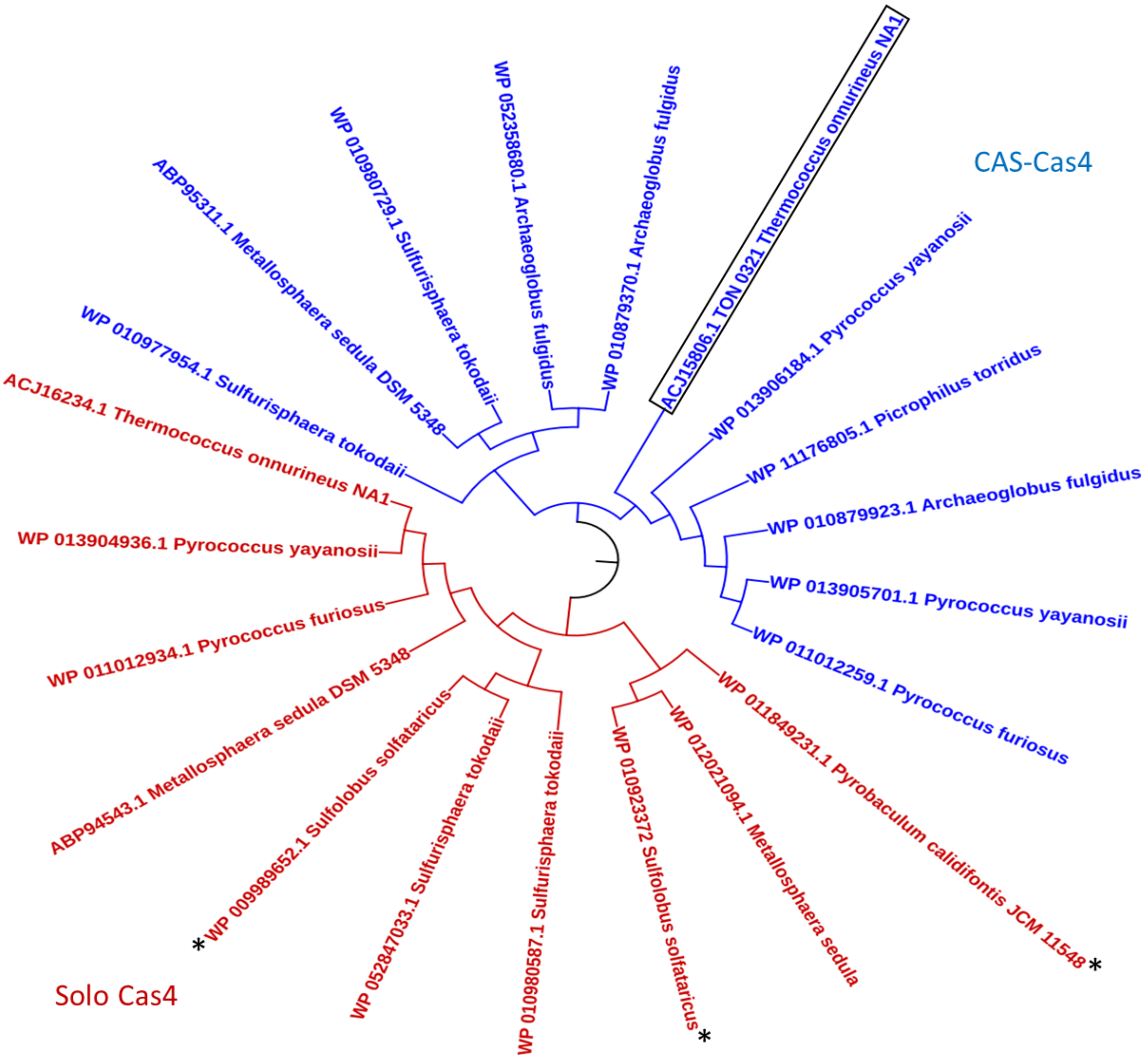


**Figure S2: Unrooted bootstrapped phylogenetic tree of Cas4 proteins.** The phylogenetic tree of Cas4 proteins from multiple archaeal species depicting the categorization of Cas4 proteins into two clear clades: Solo Cas4 proteins (Red) and CAS-Cas4 proteins (Blue). An asterisk marks the previously characterized Cas4 proteins and the protein under study (TON_0321 from *Thermococcus onnurineus*) is highlighted by a black box. The phylogenetic tree was generated using MEGA and was visualized using iTOL.

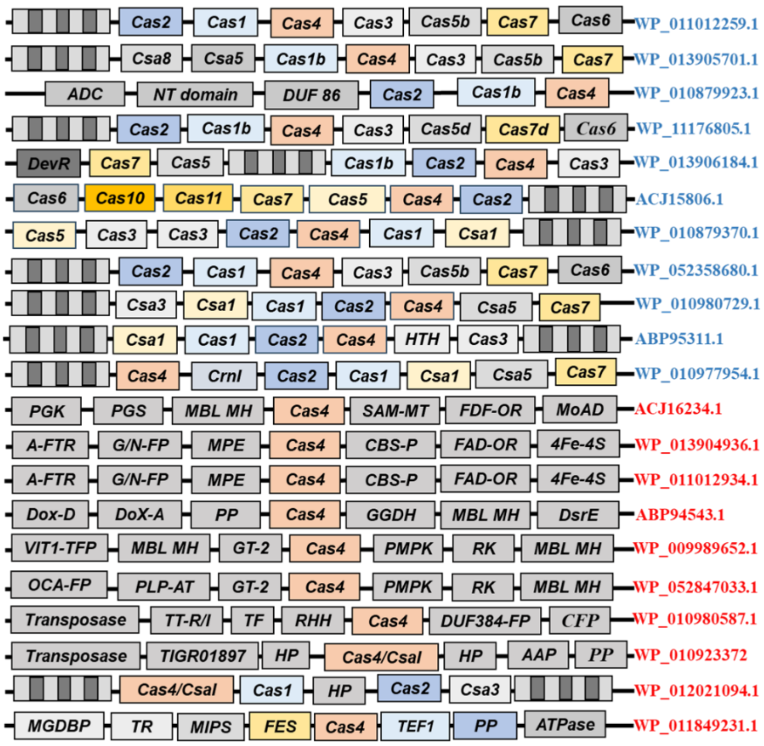


**Figure S3. A schematic of Gene cassettes.** Genes flanking the Cas4 gene used in the phylogenetic tree shown in Fig. 1. The CRISPR-associated Cas4 genes are labeled in blue, and the solo cas4 genes are labeled in red color. *PMPK*: Phosphomethyl pyrimidine kinase; *RK*: Ribokinase; *MBL MH*: MBL fold metallohydrolase; *GAT*: Class II glutamine amidotransferase; *IS605*: Transposase IS605 TnpB family; *DA*: Deacetylase; *XAP*: X-pro amino peptidase; *MGDBP*: Molybdopterin-guanine dinucleotide biosynthesis protein B; *TR*: Transcriptional regulator PadR family; *MIPS*: Myo-inositol-1-phosphate synthase; *FES*: Fe-S binding protein; *TEF1*: translation elongation factor 1A GTP binding domain family; *PP*: V-type H(+)-translocating pyrophosphatase; *ATPase*: V-type ATPase, *ADC*: Arginine decarboxylase, *NT domain*: nucleotidyltransferase domain containing protein, *DUF 86*: DUF86 containing protein, *PGK*: 2-phosphoglycerate kinase, *PGS*: 2,3-phospoglycerate synthetase, *SAM-MT*: ClassI SAM dependent methyltransferase family protein, *FDF-OR*: tungsten-containing formaldehyde ferredoxin oxidoreductase, *MoAD*: MoAD/THIS family protein, *A-FTR*: ArsR family transcriptional regulator, *G/N-FP*: Gar1/Naf1 Family protein, *MPE*: Metallophosphoesterase, *CBS-P*: CBS domain-containing protein, *FAD-OR*: FAD-dependent oxidoreductase, *4Fe-4S*: 4Fe-4S binding protein, *Dox-D*: thiosulfate dehydrogenase (quinone) subunit DoxD, *Dox-A*: thiosulfate dehydrogenase (quinone) subunit DoxA, *PP*: periplasmic protein, *GGDH*: L-glutamate gamma-semialdehyde dehydrogenase, *DsrE*: DsrE family protein, *VIT1-TFP*: VIT1/CCC1 transporter family protein, *GT-2*: Glycosyltransferase family 2 protein, *OCA-FP*: Ornithine cyclodeaminase family protein, *PLP-AT*: PLP-dependent aminotransferase family protein, *TT-R/I*: Tyrosine type recombinase/integrase, *TF*: Transcription factor, *RHH*: Ribbon helix-helix domain containing protein, *DUF384-FP*: DUF3834 domain containing protein, *CFP*: creatininase family protein, *TIGR01897*: TIGR01897 family protein, *HP*: Hypothetical protein, *AAP*: clan AA Aspartic protease.

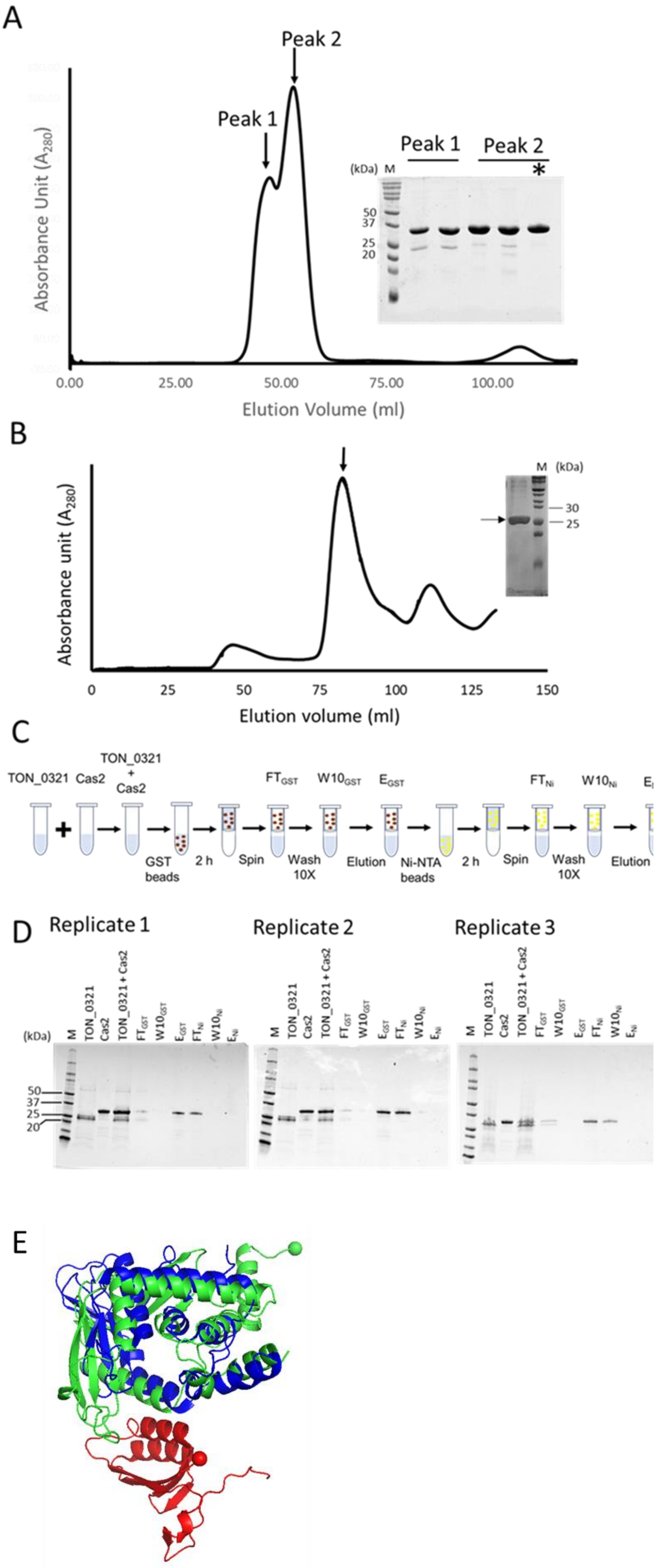
A
Peak 2
Peak 1
Absorbance Unit ($A_{280}$)
Peak 1
Peak 2
(kDa)
M
*
50
37
25
20
0.00
25.00
50.00
75.00
100.00
Elution Volume (ml)
B
Absorbance unit ($A_{280}$)
M
(kDa)
30
25
0
25
50
75
100
125
150
Elution volume (ml)
C
TON_0321
Cas2
TON_0321 + Cas2
$FT_{GST}$
$W10_{GST}$
$E_{GST}$
$FT_{Ni}$
$W10_{Ni}$
GST beads
2 h
Spin
Wash 10X
Elution
Ni-NTA beads
2 h
Spin
Wash 10X
Elution
D
Replicate 1
Replicate 2
Replicate 3
(kDa)
M
TON_0321
Cas2
TON_0321 + Cas2
$FT_{GST}$
$W10_{GST}$
$E_{GST}$
$FT_{Ni}$
$W10_{Ni}$
$E_{Ni}$
50
37
25
20
E

**Figure S4. Interaction of TON_0321 with Cas2 from *Thermococcus onnurineus*. (A)** Purification of Cas2 protein from *T. onnurineus*. The Cas2 protein was purified using size exclusion chromatography on a HiLoad 16/600 Superdex 200 column, resulting in two peaks, labeled peak 1 and peak 2. SDS-PAGE analysis was performed to assess the purity of the protein in each peak. The asterisk indicates the fraction from peak 1, free of impurities, which was used in pull-down assays to investigate the interaction between Cas2 and TON_0321 from *T. onnurineus*. (B) Purification of TON_0321 (Cas4) protein from *T. onnurineus.* The Cas4 protein was purified using size exclusion chromatography on a HiLoad 16/600 Superdex 200 column. SDS-PAGE analysis was performed to assess the purity of the protein in each peak. This same purification profile of TON_0321 has been used in Fig. 3. (C) Schematic of double pull-down assay to study Cas2 and TON_0321 interaction. The labeled steps include $FT_{GST}$ (flow-through after GST bead binding), $W10_{GST}$ (tenth wash of GST beads), $E_{GST}$ (elution from GST beads), $FT_{Ni}$ (flow-through after Ni-NTA bead binding), $W10_{Ni}$ (tenth wash of Ni-NTA beads), and $E_{Ni}$ (elution from Ni-NTA beads). These steps represent the sequential binding, washing, and elution processes used to investigate the interaction between Cas2 and TON_0321. (D) The samples were run on Bio-Rad 4-20% Mini-PROTEAN® TGX™ Precast Protein gels. The gels were stained with Coomassie Brilliant Blue. M represents marker. Lanes marked as Cas2 and TON_0321 represent individual proteins before mixing. TON_0321+ Cas2 represents the mixture of TON_0321 and Cas2 proteins. Red and yellow dots represent GST beads and Ni-NTA beads. (E) Superimposition of TON_0321 (Green) over a part of PDB 7MI4 depicting the interface between Cas4 (Blue) and Cas2 (Red) from *Geobacter sulfurreducens*. The N-terminii of TON_0321 and Cas2 are shown as green and red spheres respectively.

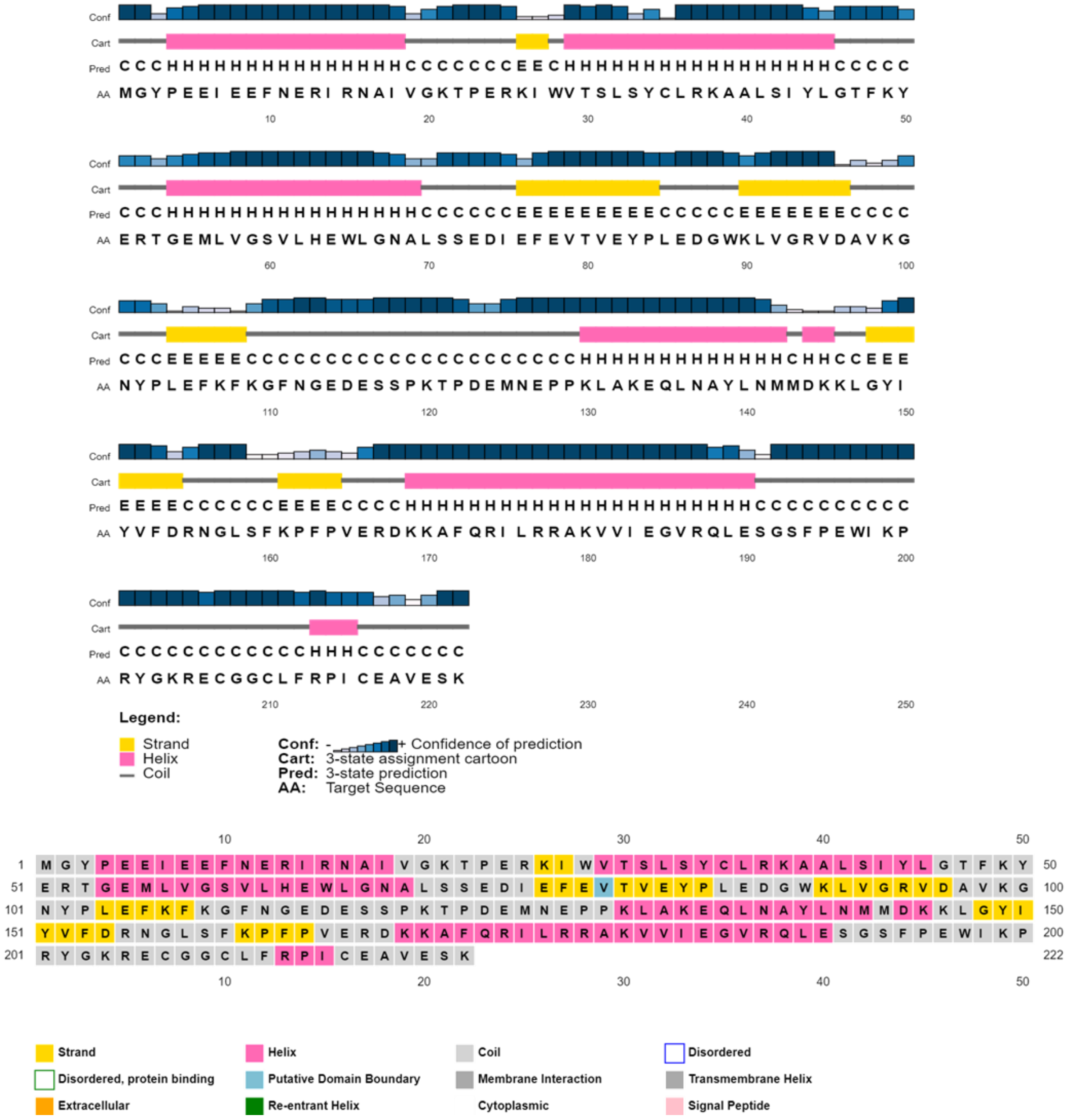


**Figure S5. *In silico* analysis of TON_0321 protein sequence.** Secondary structure (upper panel) and disorder prediction (lower panel) for TON_0321 protein using the Psipred server (75, 76).

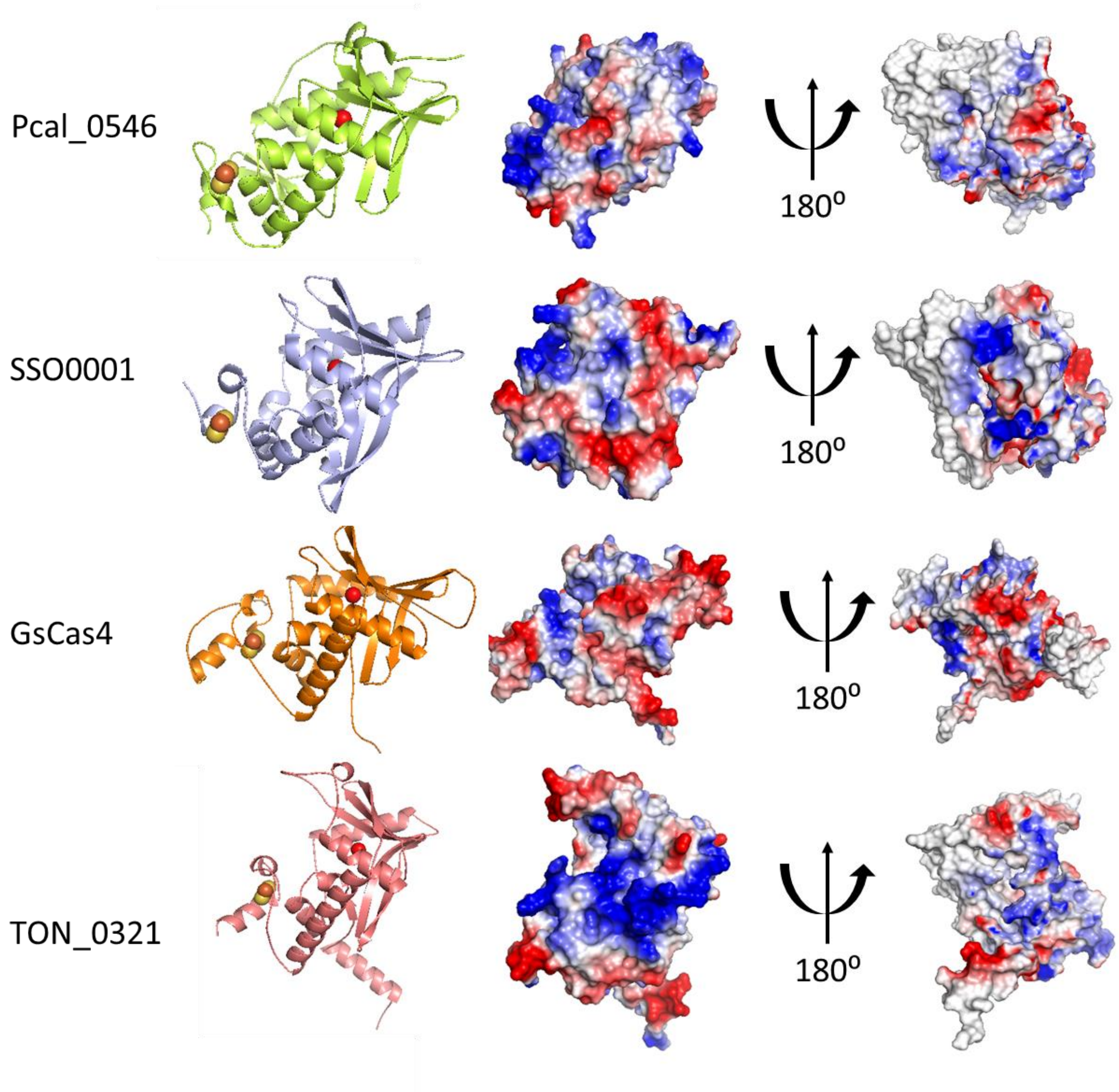


**Figure S6. The structure and surface charge distribution on the Cas4 proteins.** PDB: 4R5Q (Pcal_0546 protein from *Pyrobaculum calidifontis)*; PDB: 4IC1 (SSO0001 protein from *Sulfolobus solfataricus);* PDB: 7MI4 (GsCas4 protein from *Geobacter sulfurreducens);* TON_0321 (protein from *Thermococcus onnurineus NA1).* The first column shows the ribbon diagram. Second and third columns show surface charge distribution. Fe-S clusters are shown by orange and yellow spheres and the metal ion Magnesium by red sphere.

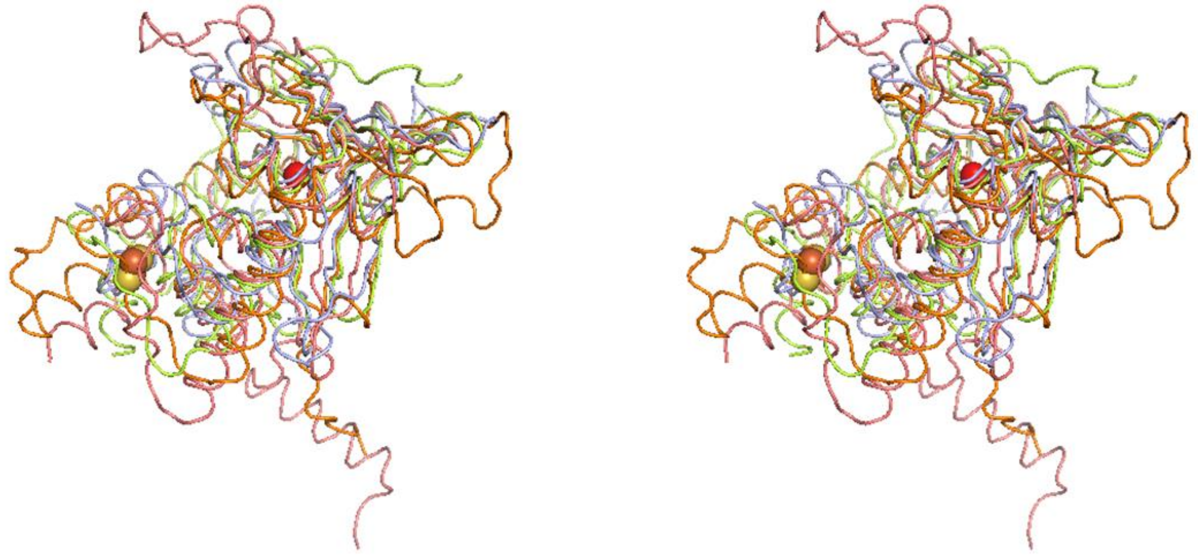

**Figure S7. Structural comparison of Cas4 proteins in stereo view.** Superimposition of Pcal_0546 (green; PDB: 4R5Q), SSO0001 (blue; PDB: 4IC1), GsCas4(orange; PDB: 7MI4), and TON_0321 (red; model). RMSD value of superimposition: TON_0321 and 4R5Q (2.53 Å), TON_0321 and SSO0001 (3.82 Å), TON_0321 and GsCas4 (4.36 Å), 4R5Q and SSO0001 (1.43 Å), GsCas4 and SSO0001 (4.41 Å), GsCas4 and 4R5Q (2.48 Å). Yellow-orange spheres denote the Fe-S cluster and the red sphere denotes $Mg^{2+}$. Despite a fairly conserved core region, the overall RMSD values are high in certain instances due to significant differences in the conformation and lengths of various loops.

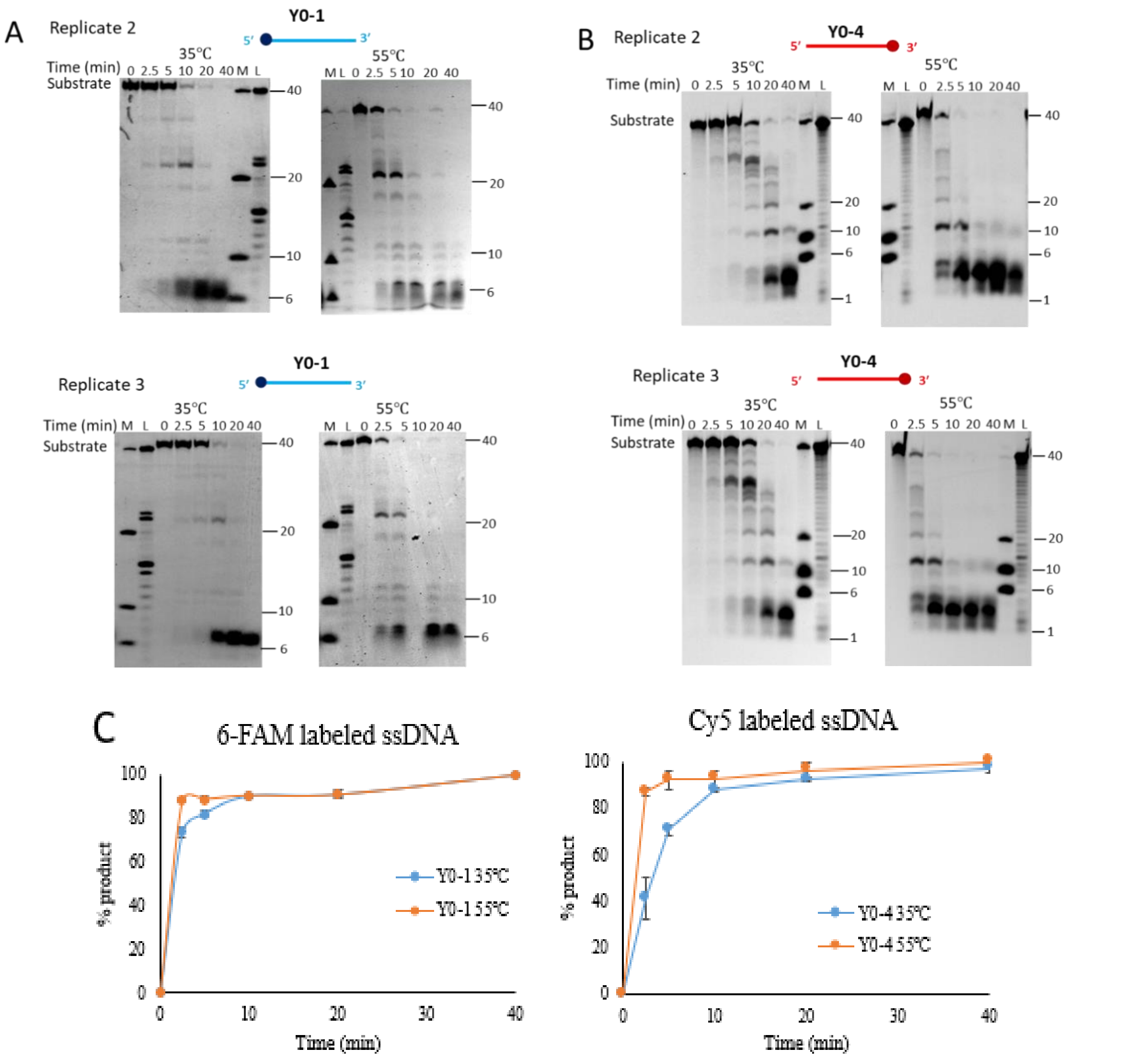


**Figure S8: Catalytic activity of TON_0321 protein on single-stranded DNA (ssDNA).** Replicates of the activity assay of TON_0321 with ssDNA are reported in Fig. 2A. The activity of TON_0321 on ssDNA labeled at 5′ end with 6-FAM (Y0-1) **(A)** and ssDNA labeled with Cy5 at 3′ end (Y0-4) **(B)**. The reaction products were resolved on an 18 % TBE-Urea PAGE. Panel (A) gels were scanned for the 6-FAM signal, and panel (B) gels were scanned for the Cy5 signal. M represents a marker made from mixing synthetic oligonucleotides of different sizes (40, 20, 10, and 6 nucleotides), and L represents a ladder made from DNase digestion of the 40 mer substrate. For replicate 2 in panel B, samples for 35°C and 55°C were run on the same gel with a common ladder and marker. **(C)** Quantitation of the product after catalytic activity of TON_0321 protein on Y0-1 and Y0-4 at 35°C and 55°C. The left Panel shows the quantitation of the product formed from activity on 6-FAM labeled ssDNA, Y0-1, and the right-side panel shows the quantitation of the product formed after activity with Cy5 labeled ssDNA, Y0-4.

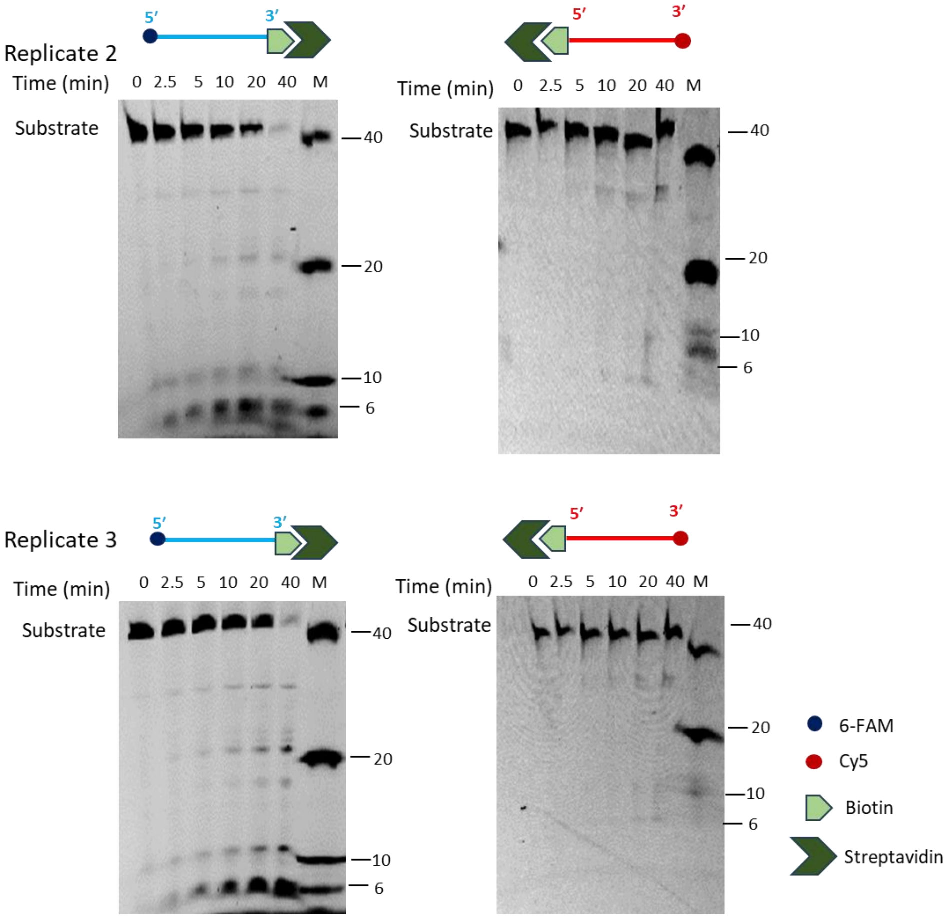


**Figure S9: Catalytic activity of TON_0321 protein on single-stranded DNA (ssDNA) with one end blocked.** The activity of TON_0321 on ssDNA labeled at 5′ end with 6-FAM, with biotin at 3′ end (Y0-1_biotin), and ssDNA labeled with Cy5 at 3′ end, and with biotin at 5′ end (Y0-4_biotin). The biotinylated oligos were incubated with 2.5 times molar excess of streptavidin to completely block the biotinylated end of the oligo. The biotin-streptavidin conjugated oligos were used as substrates for activity assay. The reaction products were resolved on an 18 % TBE-Urea PAGE. Gels were scanned for 6-FAM and Cy5 signals. M represents a marker made from mixing synthetic oligonucleotides of different sizes (40, 20, 10, and 6 nucleotides). For replicates 1 and 3, samples for Y0-1_biotin were run on the same gel with a common marker. The experiment was done in triplicates, and two replicates are shown in this figure. Replicate 1 is shown in Fig. 2A.

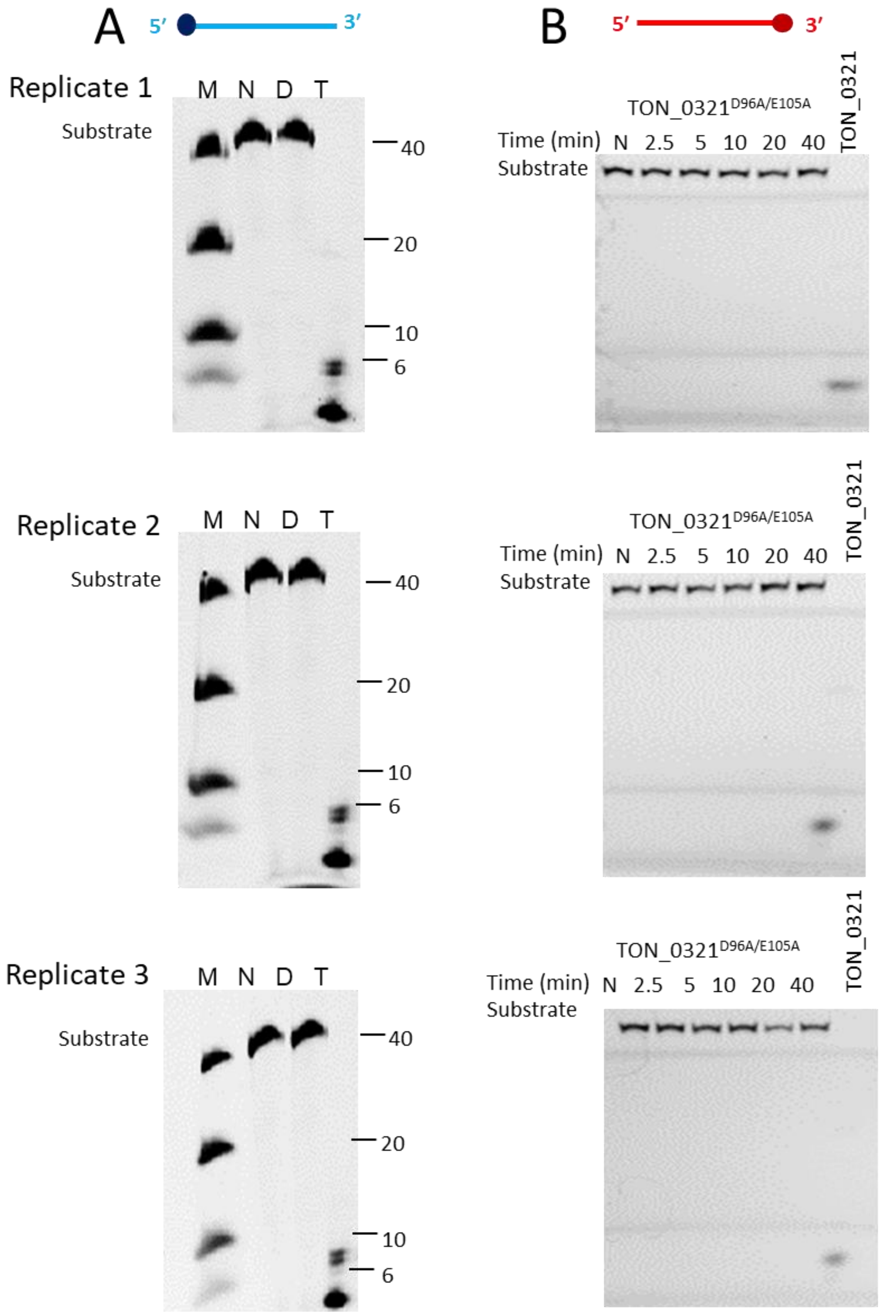


**Figure S10: Catalytic activity of wild-type TON_0321 protein and mutant protein TON_0321$^{D96A/E105A}$ on single-stranded DNA (ssDNA).** The activity of wild-type TON_0321 protein and mutant protein TON_0321$^{D96A/E105A}$ on **(A)** ssDNA labeled at 5′ end labeled with 6-FAM (Y0-1) and **(B)** ssDNA labeled with Cy5 at 3′ end (Y0-4) at 35 °C for 40 minutes. The reaction products were resolved on an 18 % TBE-Urea PAGE. Gels were scanned for 6-FAM and Cy5 signals. N: no protein control, D: mutant protein TON_0321$^{D96A/E105A}$, T: wild type TON_0321 protein. M represents a marker made from mixing synthetic oligonucleotides of different sizes (40, 20, 10, and 6 nucleotides). The experiment was done in triplicates, and three replicates are shown in the figure.

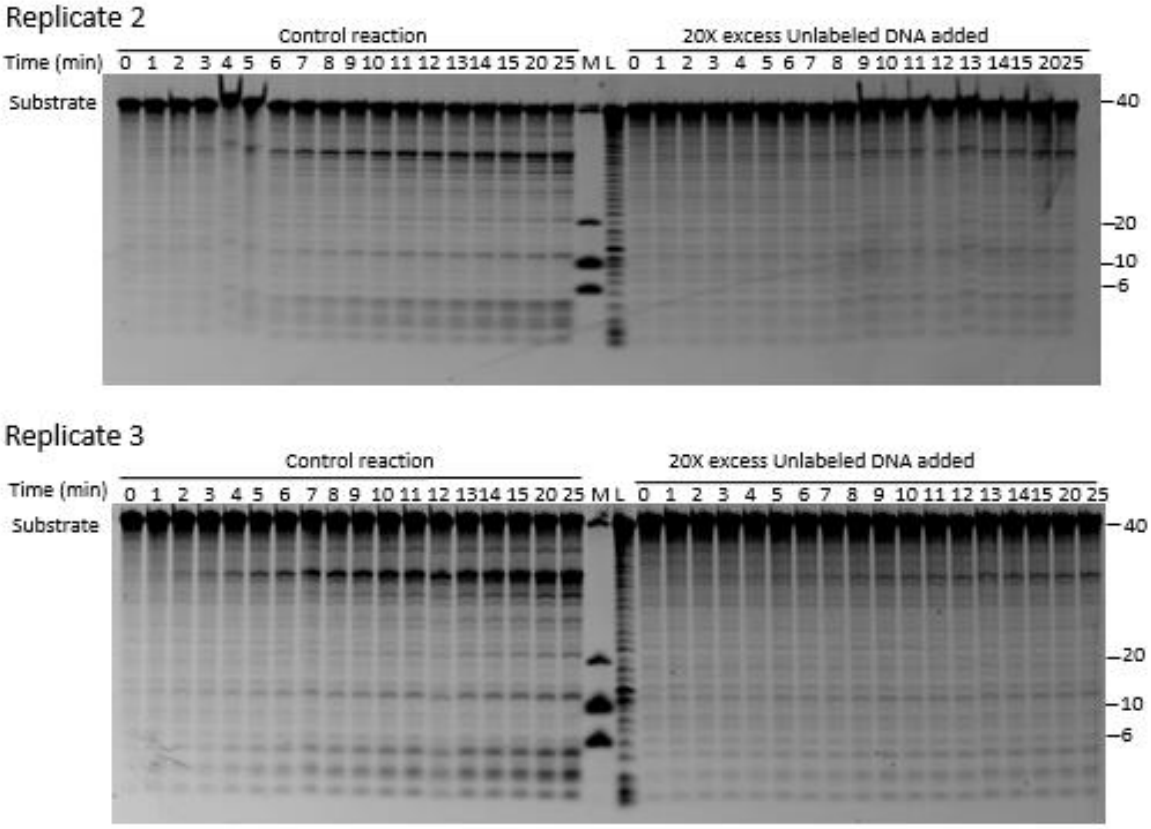


**Figure S11: TON_0321 acts in a distributive manner.** Replicates of the activity assay of TON_0321 are reported in Fig. 2C. An activity assay reaction was set with 3′ Cy5 labeled ssDNA, and the formation of products was observed temporally. In a parallel reaction, 20-fold excess unlabeled DNA of the same sequence was added after two minutes of time point. The reaction products were resolved on an 18 % TBE-Urea PAGE. The gels were scanned for Cy5 signal. M represents a marker made from mixing synthetic oligonucleotides of different sizes (40, 20, 10, and 6 nucleotides), and L represents a ladder made from DNase digestion of the 40 mer substrate.

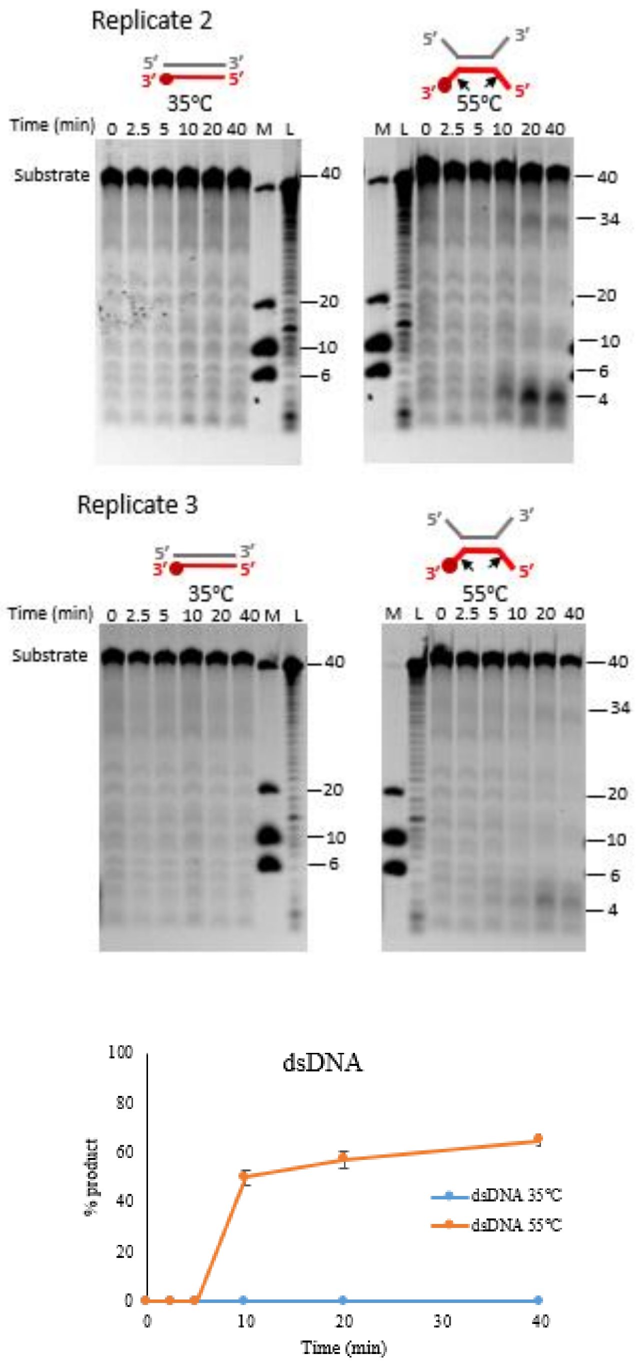


**Figure S12: Catalytic activity of TON_0321 on dsDNA.** Replicates of the activity assay of TON_0321 on dsDNA are reported in Fig. 3A. The activity of TON_0321 on dsDNA with one strand labeled with Cy5 at the 3′ end at two different temperatures, 35 °C, and 55 °C. The reaction products were resolved on an 18 % TBE-Urea PAGE. The gels were scanned for Cy5 signal. M represents a marker made from mixing synthetic oligonucleotides of different sizes (40, 20, 10, and 6 nucleotides), and L represents a ladder made from DNase digestion of the 40 mer substrate. The major cleavage site in the schematic of the DNA substrate is marked by a solid arrow. For replicate 2, samples for 35°C and 55°C were run on the same gel with a common ladder and marker. Quantitation of product after catalytic activity of TON_0321 protein on blunt-ended dsDNA substrate at 35 °C and 55 °C.

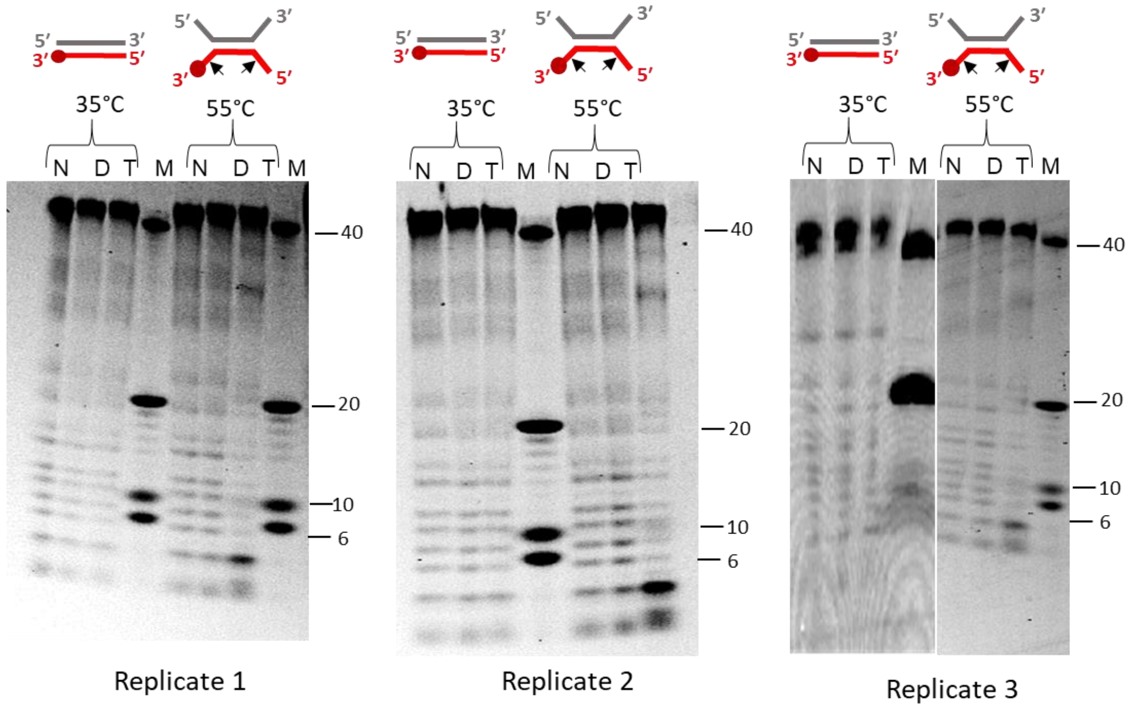


**Figure S13: Catalytic activity of wild-type TON_0321 protein and mutant protein TON_0321$^{D96A/E105A}$ on double-stranded DNA (dsDNA).** The activity of wild-type TON_0321 protein and mutant protein TON_0321$^{D96A/E105A}$ on dsDNA with one strand labeled with Cy5 at the 3′ end at two different temperatures, 35 °C and 55 °C for 40 minutes. The reaction products were resolved on an 18 % TBE-Urea PAGE. The gels were scanned for Cy5 signal. N: no protein control, D: mutant protein TON_0321$^{D96A/E105A}$, T: wild type TON_0321 protein. M represents a marker made from mixing synthetic oligonucleotides of different sizes (40, 20, 10, and 6 nucleotides). The major cleavage site in the schematic of the DNA substrate is marked by a solid arrow. The experiment was done in triplicates, and three replicates are shown in the figure.

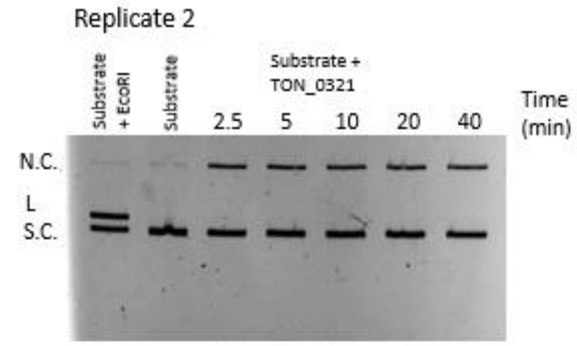


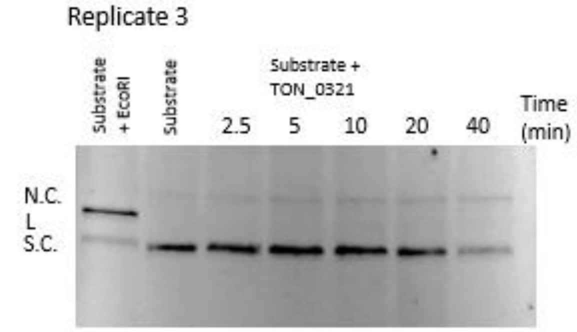


**Figure S14: Activity assay of TON_0321 using the double-stranded plasmid pIRbke8$^{mut}$.** Replicates of the activity assay of TON_0321 are reported in Fig. 3C. Ethidium Bromide stained 0.8% agarose gel showing results of cruciform assay carried out with TON_0321 protein. The lane with EcoRI represents positive control, and the substrate alone represents negative control. S.C.: supercoiled plasmid DNA, N.C.: nicked circular DNA, and L: linear DNA.

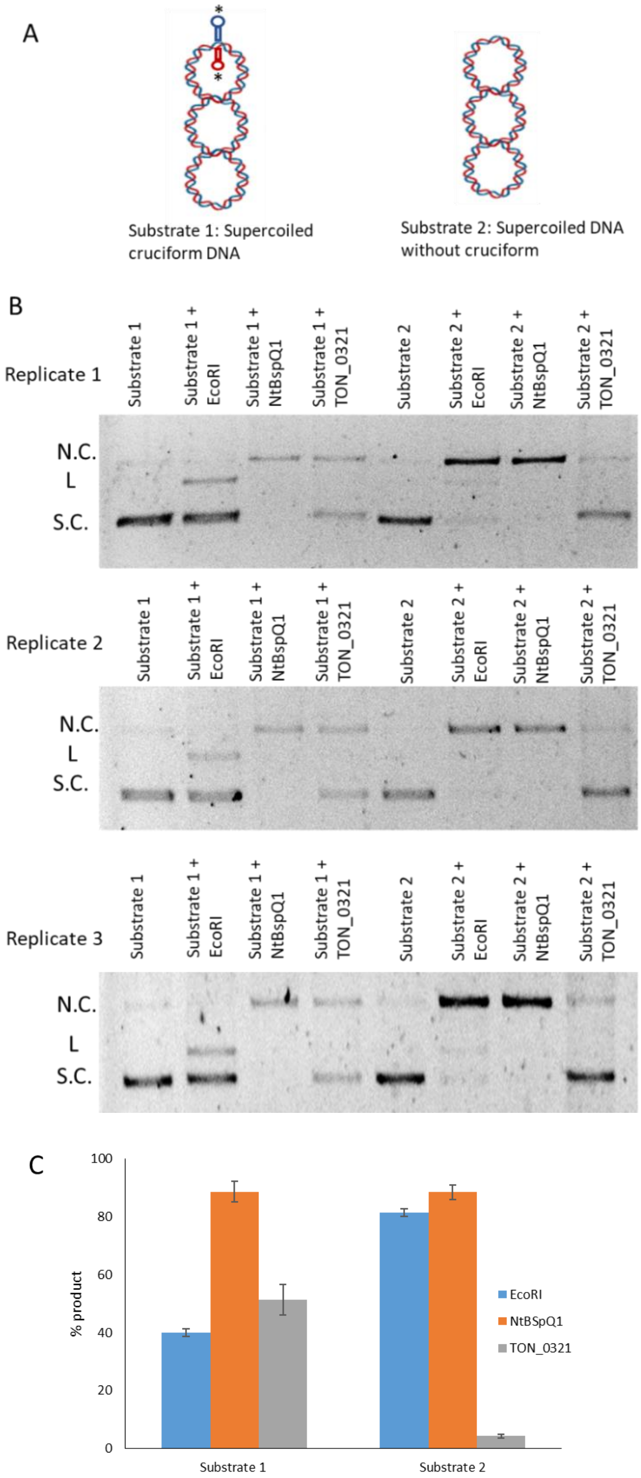


**Figure S15: Activity assay with the double-stranded plasmid pIRbke8$^{mut}$ and the modified plasmid pIRbke8$^{mut}$.** (A) The plasmid pIRbke8$^{mut}$ (substrate 1) was modified through site-directed mutagenesis to eliminate its cruciform structure, generating a variant plasmid without the cruciform (substrate 2). (B) A 0.8% agarose gel stained with ethidium bromide shows the results of an activity assay using substrate 1 and substrate 2 with the protein TON_0321, alongside controls such as EcoRI and Nt.BspQ1. The assay was conducted at 35 °C for 40 minutes. The gel displays supercoiled plasmid DNA (S.C.), nicked circular DNA (N.C.), and linear DNA (L). The experiment was performed in triplicate. (C) The quantitation from the three replicates. % product is a measure of linear plasmid in the case of reaction with EcoR1 and nicked circular plasmid in the cases of Nt.BspQ1 and TON_0321.

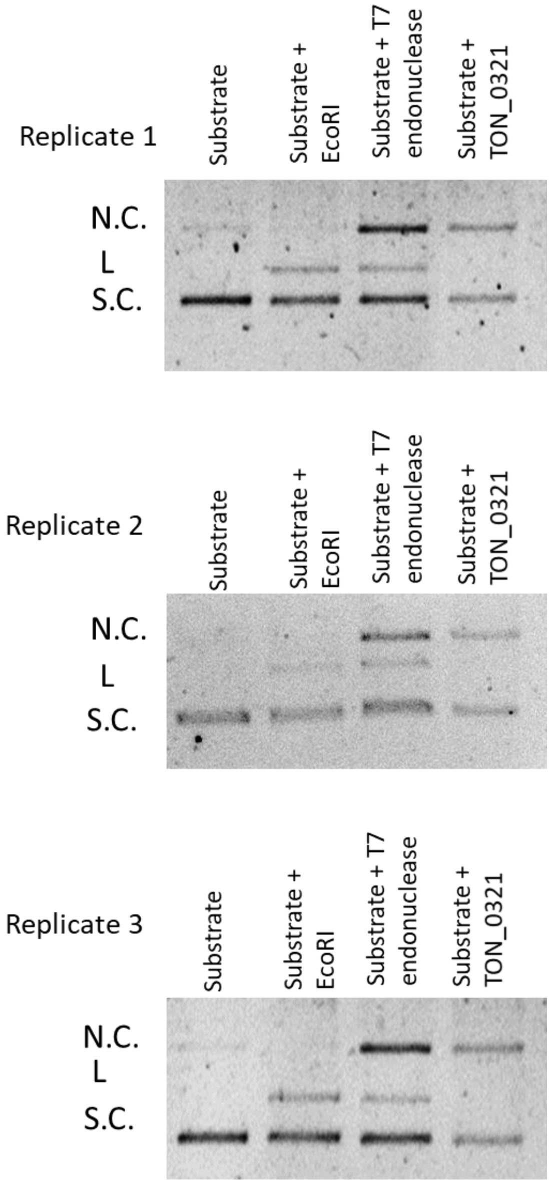


**Figure S16: Activity assay with the double-stranded plasmid pIRbke8$^{mut}$.** The plasmid pIRbke8$^{mut}$ (substrate 1) was used as a substrate for the enzymes EcoRI, T7 endonuclease I, and TON_0321. The reaction products were analyzed on a 0.8% agarose gel stained with ethidium bromide. The cruciform assay was conducted at 35 °C for 40 minutes. The gel showed supercoiled plasmid DNA (S.C.), nicked circular DNA (N.C.), and linear DNA (L). The experiment was performed in triplicate.

A

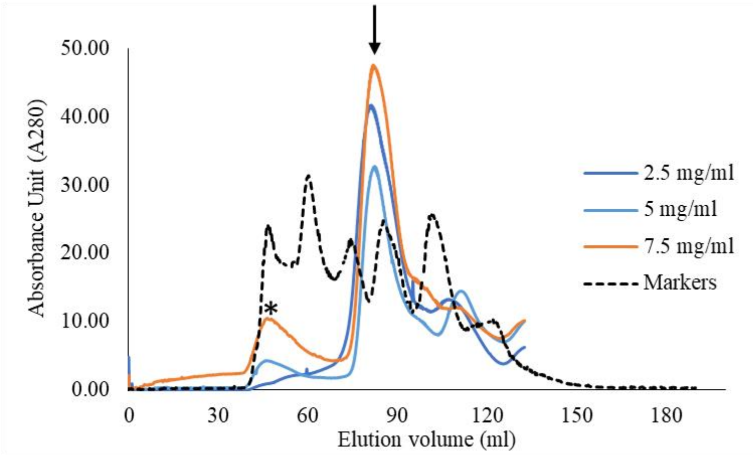


B

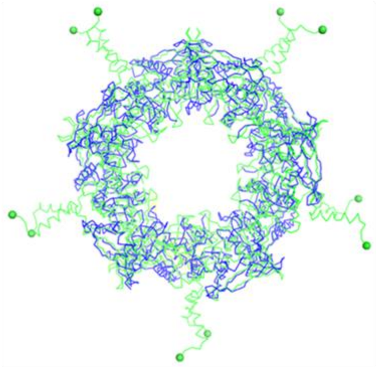

C

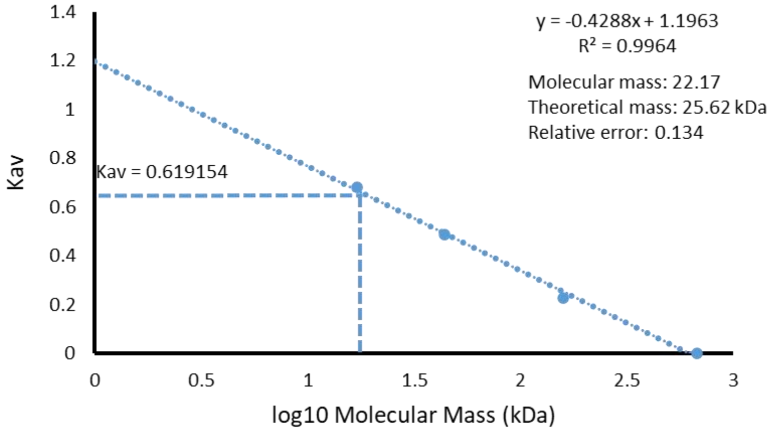


**Figure S17: Oligomeric State of TON_0321 protein.** (A) Size exclusion profile of TON_0321 under different concentrations of protein. Ton_0321 is primarily in monomeric form (peak depicted by an arrow). Asterisk depict void volume. The chromatograms for markers and protein concentration 2.5 mg/ml are same as presented in Fig. 3. (B) Superimposition of TON_0321 model (green) on SSO0001 of *Sulfolobus solfataricus* P2 (PDB: 4IC1) depicting N-ter region of TON_0321 is directed away from the interface involved in oligomerization. (C) Estimation of the oligomeric mass of TON_0321 from a standard curve generated using gel filtration markers (Vitamin B12, Myoglobin, Ovalbumin and gamma globulin). Kav was calculated as (Ve-Vo)/(Vt-Vo) where Ve, Vo, and Vt are elution volume, void volume, and total volume, respectively. Thyroglobulin (670 kDa) was used to determine the column's void volume.

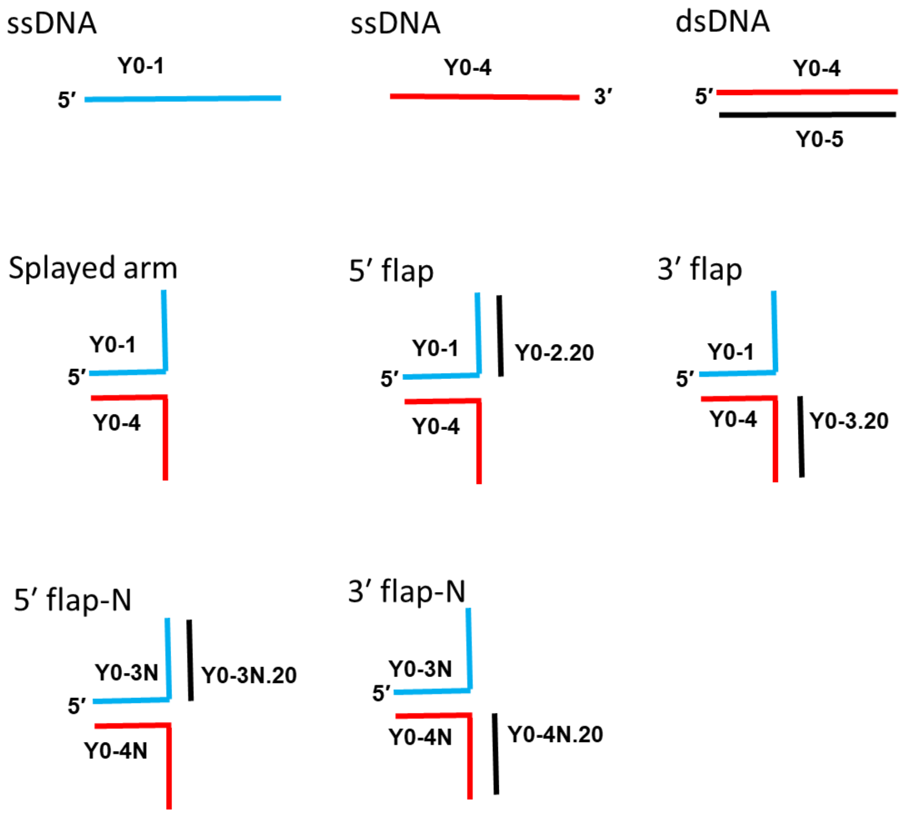


**Figure S18. Schematic representation of substrates used in nuclease assays.** Cartoon representation of the substrates used in activity assays. Blue represents 6-FAM and red represents Cy5 labeled strands. 5' flap-N and 3' flap-N has sequences flipped at the branching points with respect to 5' flap and 3' flap substrates. The sequence details are available in Table S1.

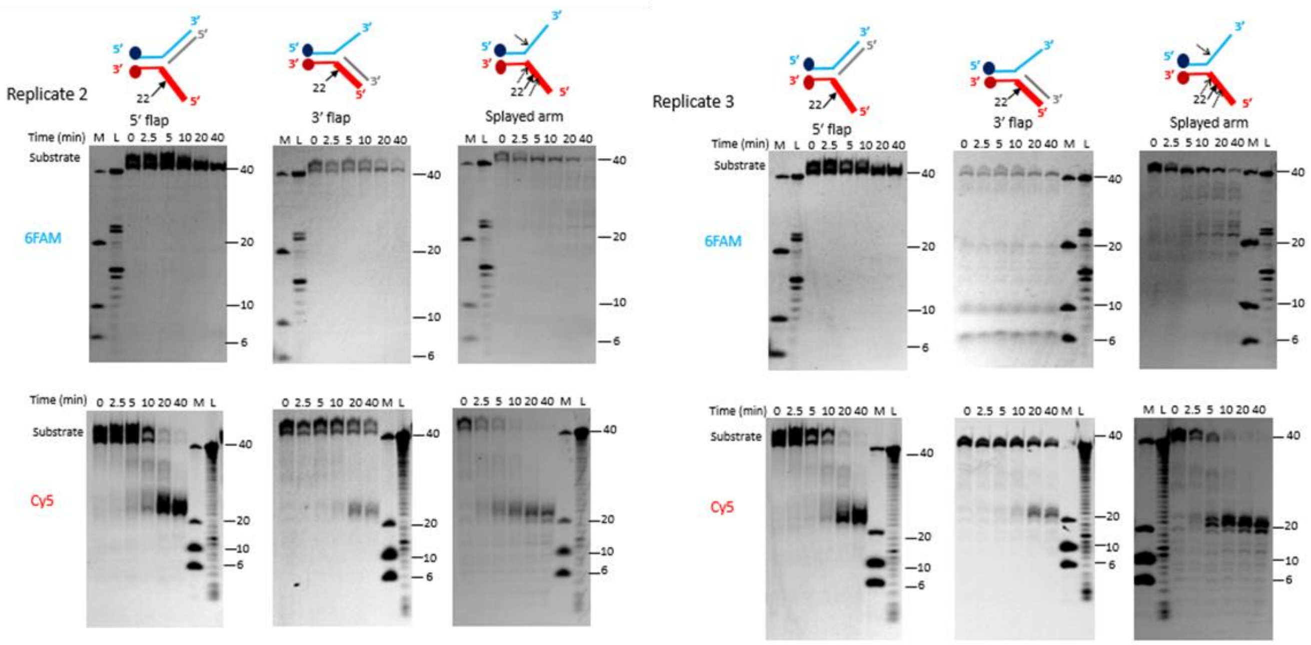


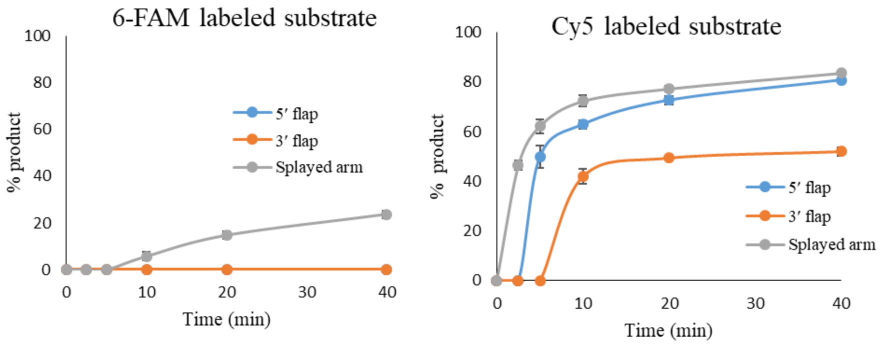


**Figure S19: Catalytic activity of TON_0321 on branched DNA molecules.** Replicates of the activity assay of TON_0321 on different branched DNA substrates, 5′ flap, 3′ flap, and splayed arm are reported in Fig. 4A. The catalytic activity of TON_0321 protein was checked on branched DNA molecules: 5′ flap, 3′ flap and splayed arm. Each substrate has two labels: one strand labeled at 5′ end with 6-FAM and another strand labeled at 3′ end with Cy5. The reaction products were resolved on an 18 % TBE-Urea PAGE. The same gel was scanned for the 6-FAM signal (upper panel of both replicates) and for the Cy5 signal (lower panel of both replicates). M represents a marker made from mixing synthetic oligonucleotides of different sizes (40, 20, 10, and 6 nucleotides), and L represents a ladder made from DNase digestion of the 40 mer substrate. The major cleavage site in the schematic of the DNA substrate is marked by a solid arrow. For replicate 2, 5′ flap and 3′ flap samples were run on the same gel sharing a common ladder and marker for Cy5 signal. For replicate 2, 3′ flap and splayed arm samples were run on the same gel sharing a common ladder and marker for 6-FAM signal. Quantitation of product after catalytic activity of TON_0321 protein on 5′ flap, 3′ flap, and splayed arm at 35°C. The left Panel shows the quantitation for the 6-FAM signal, and the right panel shows the quantitation for the Cy5 signal.

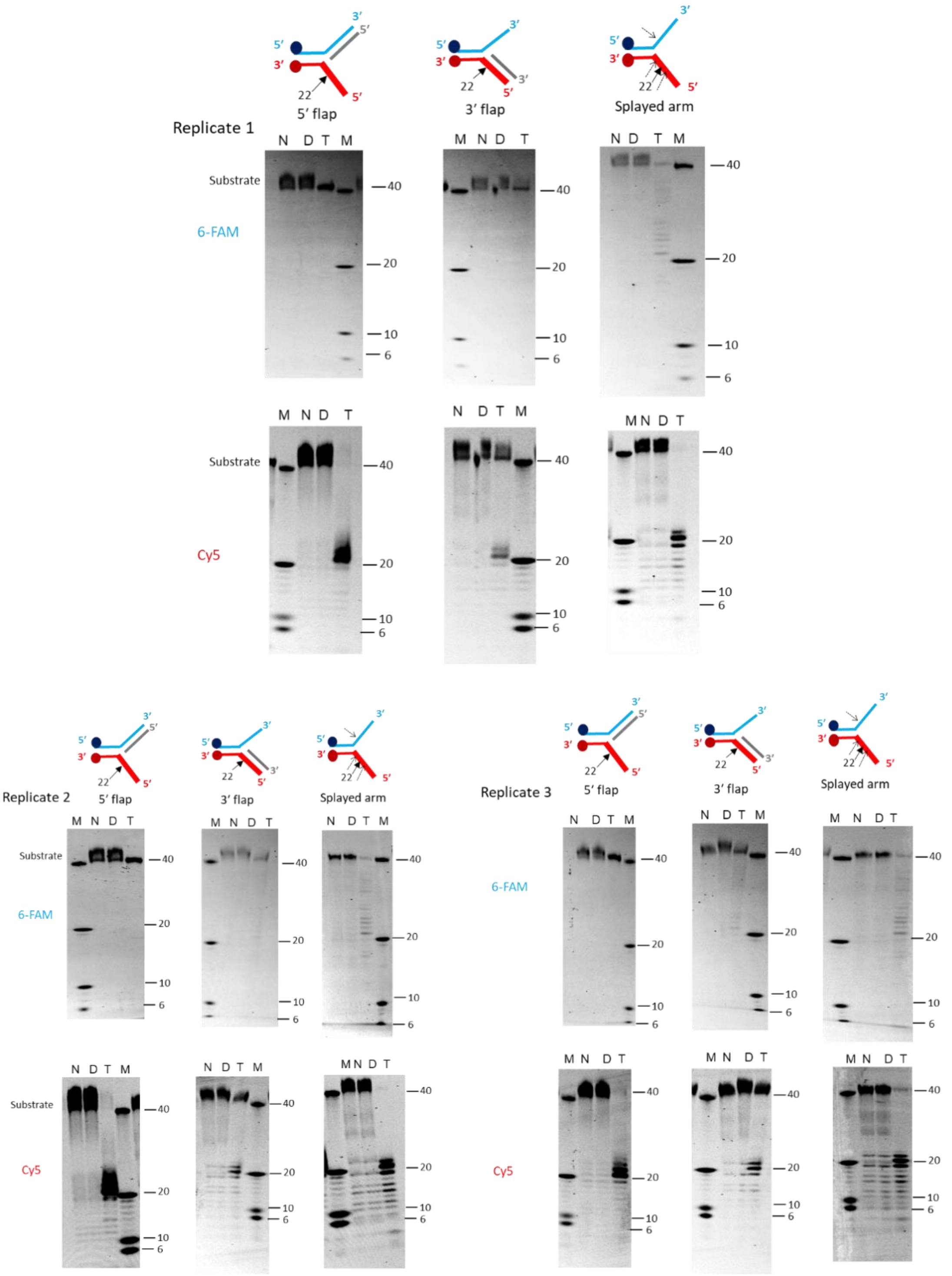

Replicate 1
5′ flap
3′ flap
Splayed arm
Substrate
6-FAM
Cy5
N D T M
M N D T
40
20
10
6
Replicate 2
Replicate 3

**Figure S20: Catalytic activity of wild-type TON_0321 protein and mutant protein TON_0321$^{D96A/E105A}$ on branched DNA molecules.** The activity of wild-type TON_0321 protein and mutant protein TON_0321$^{D96A/E105A}$ on different branched DNA substrates, 5′ flap, 3′ flap, and splayed arm were carried out at 35°C for 40 minutes. Each substrate has two labels: one strand labeled at 5′ end with 6-FAM and another strand labeled at 3′ end with Cy5. The reaction products were resolved on an 18 % TBE-Urea PAGE. The same gel was scanned for the 6-FAM signal (upper panel of all replicates) and the Cy5 signal (lower panel of all replicates). N: no protein control, D: mutant protein TON_0321$^{D96A/E105A}$, T: wild type TON_0321 protein. M represents a marker made from mixing synthetic oligonucleotides of different sizes (40, 20, 10, and 6 nucleotides). For replicate 1, 5′ flap and 3′ flap samples were run on the same gel sharing a common marker for 6-FAM signal. For replicate 1, 3′ flap and splayed arm samples were run on the same gel sharing a common marker for Cy5 signal. For replicate 2, 5′ flap and splayed arm samples were run on the same gel sharing a common marker for Cy5 signal. The samples for replicate 2 of 3′ flap and replicate 3 of 5′ flap were run on the same gel sharing a common marker for 6-FAM signal. The samples for replicate 2 and replicate 3 of 3′ flap were run on the same gel sharing a common marker for Cy5 signal. The replicate 3 of 3′ flap and splayed arm samples were run on the same gel sharing a common marker for 6-FAM signal. The major cleavage site in the schematic of the DNA substrate is marked by a solid arrow. The experiment was done in triplicates, and three replicates are shown in the figure.

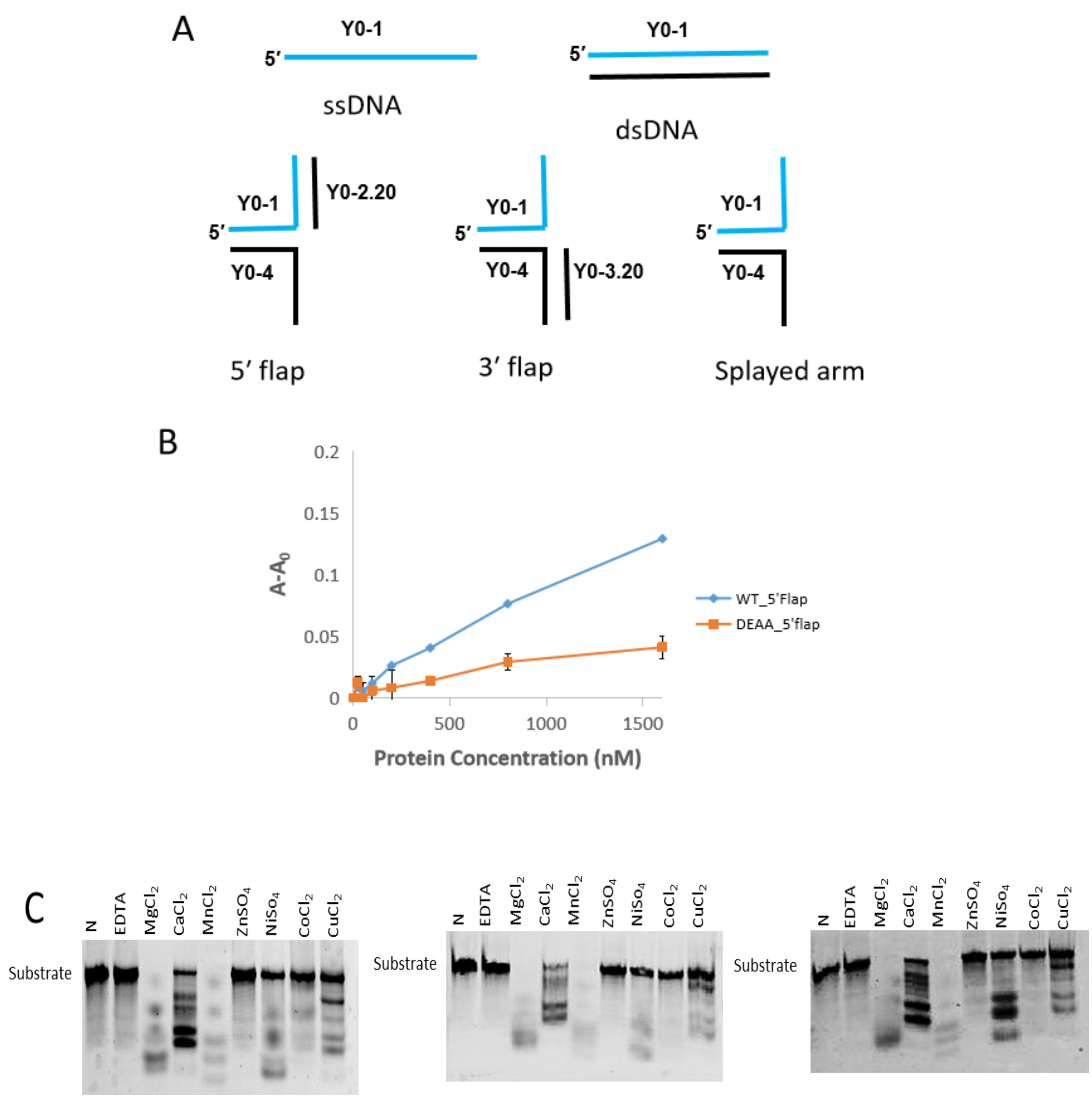


**Figure S21. (A) Schematic representation of substrates used in fluorescence anisotropy.** Cartoon representation of the substrates used in DNA substrate binding studies. Blue represents 6-FAM labeled strand. (B) Binding study of TON_0321 wild type and mutant TON_0321$^{D96A/E105A}$ protein with 6-FAM labelled 5'flap as substrate using Fluorescence anisotropy. The Y-axis shows a change in anisotropy ($A - A_0$), where A is observed anisotropy and $A_0$ is anisotropy of DNA substrate alone. (C) Activity of TON_0321 protein on ssDNA substrate in presence of different metal ions. Gel was scanned for Cy5 signal.

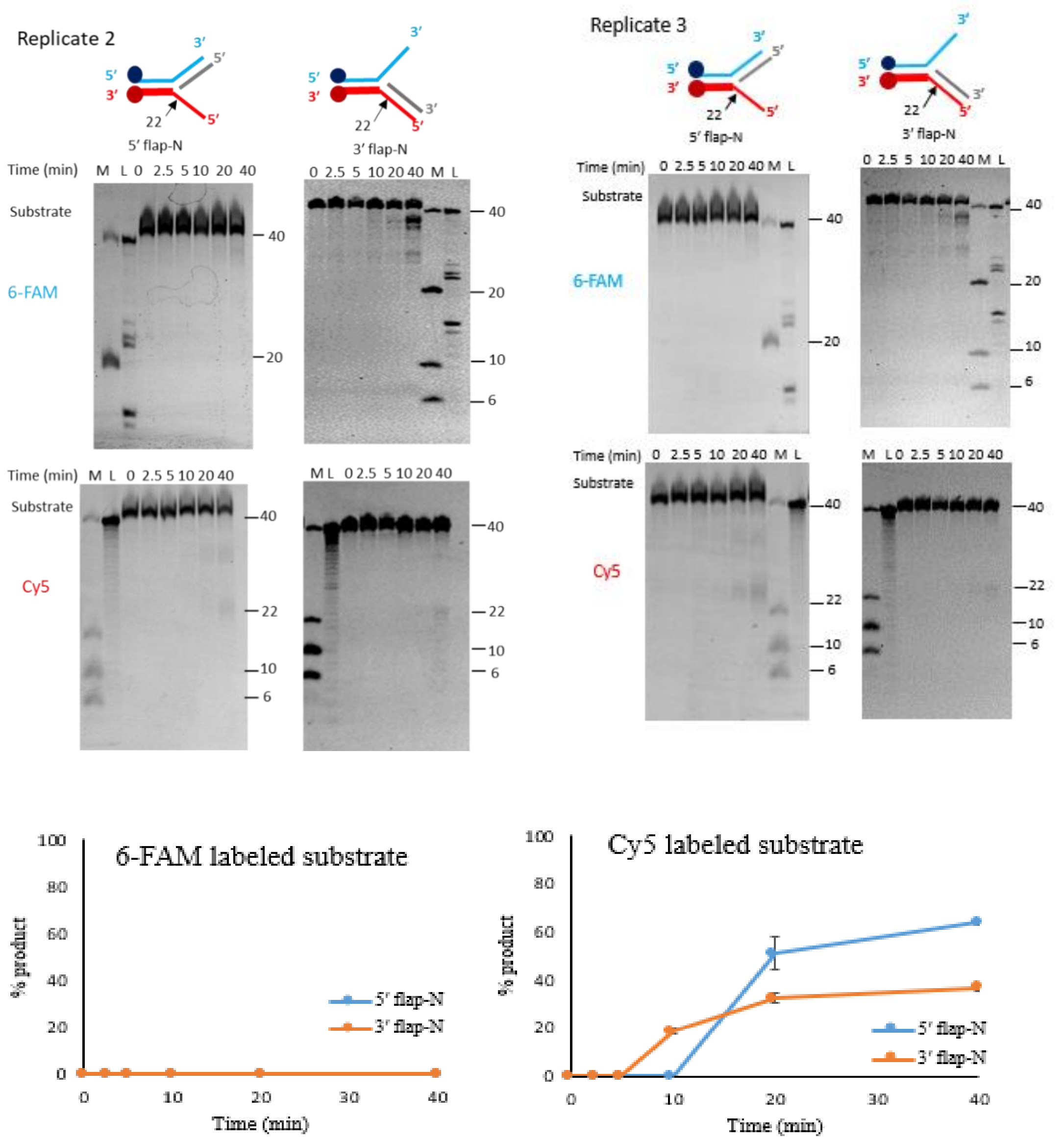


**Figure S22: TON_0321 enzyme is a secondary structure-specific endonuclease.** Replicates of the activity assay of TON_0321 on new 5′ flap and 3′ flap substrates having sequence flipped around the branch point, 5′ flap-N, and 3′ flap-N are reported in Fig. 5A and Fig. 5B. The catalytic activity of TON_0321 protein on branched DNA molecules, 5′ flap-N and 3′ flap-N with each having one strand labeled at 5′ end with 6-FAM and another strand labeled at 3′ end with Cy5. The reaction products were resolved on 18 % TBE-Urea PAGE. The gels were scanned for 6-FAM signal (upper panel) and Cy5 signal (lower panel). M represents a marker made from mixing synthetic oligonucleotides of different sizes (40, 20, 10, and 6 nucleotides), and L represents a ladder made from DNase digestion of the 40 mer substrate. The samples for replicate 2 and 3 of 5′ flap-N were run on the same gel sharing a common marker and ladder for Cy5 signal. The major cleavage site in the schematic of branched DNA substrates is marked by a solid arrow. Quantitation of product after catalytic activity of TON_0321 protein on 5′ flap, 3′ flap, and splayed arm at 35°C. The left Panel shows the quantitation for the 6-FAM signal, and the right panel shows the quantitation for the Cy5 signal.

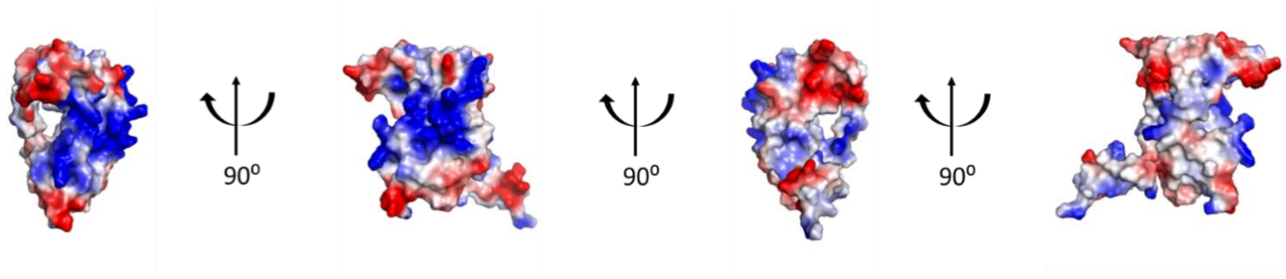


**Figure S23. Surface charge distribution on the TON_0321 protein.** TON_0321 protein has surface-exposed positively charged residues organized into a distinct patch optimum for interaction with DNA duplex.

**Table S1. Oligonucleotides used for cloning and Site-Directed Mutagenesis**

| Name | Sequence (5′ -3′) |
|---|---|
| **TON_0321_FP** | CGGGATCCATGGGCTATCCGGAAGAAATTG |
| **TON_0321_RP** | CCGCTCGAGTCATTTAGACTCAACGGCCTC |
| **ToN_0321_DEAA_FP** | GTGGCCGCAGTAAAGGGGAATTATCCACTTGCGTTCAA |
| **ToN_0321_DEAA_RP** | TTGAACGCAAGTGGATAATTCCCCTTTACTGCGGCCAC |
| **Cruciform_SDM_FP** | CACACCCGTCCTGTCTACGCCGGACGCATC |
| **Cruciform_SDM_RP** | GATGCGTCCGGCGTAGACAGGACGGGTGTG |
| **CAS 2 F.P. BamHI** | CGGGATCCATGTACGTGGTCATCGTCTACGATG |
| **CAS 2 R.P. XhoI** | CCGCTCGAGCTAAATGATGTCCTCCAGAGGGTTC |

**Table S2. Details of DNA oligonucleotides and substrates used in nuclease assays.**

**Table S2A.** Sequences of DNA oligonucleotides used in annealing synthetic substrates for activity assays

| Name | Sequence (5′ - 3′) |
|---|---|
| Y0-1 | ATTCTACCAGTGCCTTGCTA GGACATCTTTGCCCACCTGC |
| Y0-4 | ATCCTCTAGACAGCTCCATGTAGCAAGGCACTGGTAGAAT |
| Y0-5 | ATTCTACCAGTGCCTTGCTACATGGAGCTGTCTAGAGGAT |
| Y0-3N | CATGGAGCTGTCTAGAGGATGCTTGACGATTACAACAGAT |
| Y0-4N | TAGCAAGGCACTGGTAGAATATCCTCTAGACAGCTCCATG |
| Y0-2.20 | GCAGGTGGGCAAAGATGTCC |
| Y0-3.20 | CATGGAGCTGTCTAGAGGAT |
| Y0-4N.20 | ATTCTACCAGTGCCTTGCTA |
| Y0-3N.20 | ATCTGTTGTAATCGTCAAGC |
| Y0-1F | [6-FAM] ATTCTACCAGTGCCTTGCTAGACATCTTTGCCCACCTGC |
| Y0-4C | ATCCTCTAGACAGCTCCATG TAGCAAGGCACTGGTAGAAT [Cy5] |
| Y0-4N-CY5 | TAGCAAGGCACTGGTAGAATATCCTCTAGACAGCTCCATG [Cy5] |
| Y0-3N-6-FAM | [6-FAM] CATGGAGCTGTCTAGAGGATGCTTGACGATTACAACAGAT |

**Table S2B.** Combination of DNA oligonucleotides used to generate various joint DNA molecules

| Substrate | Unlabeled | Labeled |
|---|---|---|
| **ssDNA** | Y0-1 | Y0-1F |
| **ssDNA** | Y0-4 | Y0-4C |
| **dsDNA** | Y0-4+ Y0-5 | Y0-4C + Y0-5 |
| **5′ flap** | Y0-1+Y0-4+Y0-2.20 | Y0-1F +Y0-4C +Y0-2.20 |
| **3′ flap** | Y0-1+Y0-4 + Y0-3.20 | Y0-1F +Y0-4C + Y0-3.20 |
| **Splayed arm** | Y0-1+Y0-4 | Y0-1F +Y0-4C |
| **5′ flap-N** | Y0-3N+Y0-4N+Y0-2.20 | Y0-3N-6-FAM + Y0-4N-Cy5 + Y0-3N.20 |
| **3′ flap-N** | Y0-3N+Y0-4N+Y0-3.20 | Y0-3N-6-FAM + Y0-4N-Cy5 + Y0-4N.20 |

**Table S2C.** Sequences of DNA oligonucleotides used for generating DNA ladders.

| Name | Sequence (5′ - 3′) |
|---|---|
| Y0-1F | [6-FAM] ATTCTACCAGTGCCTTGCTAGGACATCTTTGCCCACCTGC |
| LO-1.20F | [6-FAM] ATTCTACCAGTGCCTTGCTA |
| LO-1.10F | [6-FAM] ATTCTACCAG |
| LO-1.05F | [6-FAM] ATTCT |
| Y0-4C | ATCCTCTAGACAGCTCCATGTAGCAAGGCACTGGTAGAAT [Cy5] |
| Y0-1.20C | TAGCAAGGCACTGGTAGAAT [Cy5] |
| Y0-1.10C | CTGGTAGAAT [Cy5] |
| Y0-1.05C | AGAAT [Cy5] |

**Table S3. Details of DNA oligonucleotides and substrates used in fluorescence anisotropy.**

**Table S3A.** Sequences of DNA oligonucleotides used in annealing synthetic substrates for fluorescence anisotropy

| Name | Sequence (5′ - 3′) |
|---|---|
| Y0-1 | ATTCTACCAGTGCCTTGCTA GGACATCTTTGCCCACCTGC |
| Y0-4 | ATCCTCTAGACAGCTCCATGTAGCAAGGCACTGGTAGAAT |
| Y0-8 | TAAGATGGTCACGGAACGATCCTGTAGAAACGGGTGGACG |
| Y0-2.20 | GCAGGTGGGCAAAGATGTCC |
| Y0-3.20 | CATGGAGCTGTCTAGAGGAT |
| Y0-1F | [6-FAM]ATTCTACCAGTGCCTTGCTAGGACATCTTTGCCCACCTG |

**Table S3B.** Combination of DNA oligonucleotides used to generate various joint DNA molecules

| Substrate | Unlabeled | Labeled |
|---|---|---|
| **ssDNA** | Y0-1 | Y0-1F |
| **dsDNA** | Y0-1+ Y0-8 | Y0-1F + Y0-8 |
| **5′flap** | Y0-1+Y0-4+Y0-2.20 | Y0-1F +Y0-4 + Y0-2.20 |
| **3′flap** | Y0-1+Y0-4 + Y0-3.20 | Y0-1F +Y0-4 + Y0-3.20 |
| **Splayed arm** | Y0-1+Y0-4 | Y0-1F +Y0-4 |

**Table S4: Raw Data for Fluorescence Anisotropy**

| Protein (nM) | ToN_0321$^{WT}$ | | | | | | | TON_0321 $^{D96A/E105A}$ |
|---|---|---|---|---|---|---|---|---|
| **Average Total Intensity:** Calculated as $(I_{\|\|} + 2I_{\perp})$, where $I_{\|\|}$ and $I_{\perp}$ are intensities in parallel and perpendicular directions | | | | | | | | |
| | 5' Flap | 3' Flap | SA | ssDNA | dsDNA | 5' Flap-N | 3' Flap-N | 5' Flap |
| 1600 | 58796.67 | 60920 | 71779.33 | 63520.33 | 69766 | 31012 | 30113.33 | 34094.33 |
| 800 | 63319.67 | 67031.33 | 61562 | 71464.33 | 78562.67 | 31616 | 29608.33 | 33335 |
| 400 | 72238.33 | 72942.33 | 64058 | 71986.33 | 76652 | 37583.67 | 28850 | 34800 |
| 200 | 71333.33 | 74655.33 | 66330 | 72828 | 73243.33 | 31491.33 | 28461.67 | 31326.67 |
| 100 | 69536.67 | 76288 | 65299 | 70853.67 | 73642.33 | 29785.33 | 28416 | 34186.67 |
| 50 | 70783.67 | 72071.33 | 65742 | 73260 | 73889.67 | 26252.67 | 28141.33 | 32012 |
| 25 | 70460.67 | 71753 | 66561 | 71989.33 | 74783 | 29657 | 27450.67 | 33029 |
| 0 | 73035.67 | 72353.33 | 69699 | 72652 | 72237.33 | 29108.33 | 29191.33 | 32534 |
| **Average Anisotropy:** Calculated as $(I_{\|\|} - I_{\perp}) / (I_{\|\|} + 2I_{\perp})$, where $I_{\|\|}$ and $I_{\perp}$ are intensities in parallel and perpendicular directions | | | | | | | | |
| Protein (nM) | ToN_0321$^{WT}$ | | | | | | | TON_0321 $^{D96A/E105A}$ |
| | 5' Flap | 3' Flap | SA | ssDNA | dsDNA | 5' Flap-N | 3' Flap-N | 5' Flap |
| 1600 | 0.155732 | 0.11581 | 0.090413 | 0.065608 | 0.084775 | 0.13289 | 0.135048 | 0.068572 |
| 800 | 0.096439 | 0.09387 | 0.083964 | 0.03371 | 0.044592 | 0.101616 | 0.082484 | 0.056711 |
| 400 | 0.058679 | 0.059222 | 0.069541 | 0.024144 | 0.029921 | 0.070636 | 0.056627 | 0.041406 |
| 200 | 0.041841 | 0.034674 | 0.04646 | 0.016601 | 0.024328 | 0.058826 | 0.036031 | 0.035711 |
| 100 | 0.031314 | 0.030784 | 0.040822 | 0.018616 | 0.029651 | 0.043633 | 0.031237 | 0.033465 |
| 50 | 0.025347 | 0.023296 | 0.036293 | 0.018821 | 0.025399 | 0.041327 | 0.024217 | 0.027762 |
| 25 | 0.031678 | 0.028367 | 0.037019 | 0.025048 | 0.027766 | 0.026476 | 0.054626 | 0.040162 |
| 0 | 0.021928 | 0.020321 | 0.023592 | 0.016533 | 0.020536 | 0.019269 | 0.024737 | 0.027524 |
